# Identification of mobile element insertion from whole genome sequencing data using deep neural network model

**DOI:** 10.1101/2023.03.07.531451

**Authors:** Xiaofei Xu, Yu Huang, Xuegang Wang, Jing Cheng, Huijun Yuan, Fengxiao Bu

## Abstract

Mobile element insertions (MEIs) are a major contributor to genome evolution and play an essential role in the regulation of gene expression, as well as being implicated in various human diseases. This study introduces DeepMEI, a tool based on a convolutional neural network model that transforms the MEI identification process into an image recognition problem and automatically learns complex and abstract representations of MEI features in whole genome sequencing data. DeepMEI outperformed existing tools in the benchmark dataset from the Genome in a Bottle consortium, with a precision of 0.90 and recall of 0.70. Moreover, factors such as sequencing depth, ME integrity, and genome mappability can affect MEI identification accuracy. Using DeepMEI, we reanalyzed 3,202 high-coverage whole-genome sequencing samples from the 1000 Genome Project (1kGP) phase 4 release, discovering 1.71-fold more non-reference MEIs, totaling 6,218,088, with 92.2% of the increase coming from rare MEIs (allele frequency <1%). This enhances our understanding of MEIs’ role in human disease and evolution. The DeepMEI tool and the updated 1kGP MEI dataset can be accessed at https://github.com/xuxif/DeepMEI.

## INTRODUCTION

Mobile elements (MEs), also known transposable elements, transposons, or jumping genes, compose up to 45% of the human genome (1, 2). MEs can be categorized as DNA transposons, long terminal repeat (LTR) retrotransposons, and non-LTR retrotransposons. Most MEs in the human genome have been silenced during evolution due to accumulated truncations and host repressive mechanisms (3) such as methylation (4), heterochromatin formation (5), and piwi-interacting RNA regulation (6). However, some structurally and functionally intact non-LTR retrotransposons, including LINE-1 (L1; ∼6000 bp in length), Alu (∼300 bp) and SINE-VNTR-Alu (SVA; ∼2000 bp) elements, still possess transposition activity (7, 8). ME-mediated transposition and homologous recombination are the main sources of genome instability and play essential roles in genome evolution (9, 10), transcription regulation (11), and three-dimensional genome organization (12, 13). MEs are ideal genetic markers for evolutionary and populational studies, given the homoplasy-free (14) and identical-by-descent (15) properties of non-autonomous transposition (16). Further, with respect to genic insertions, MEIs have the potential to disrupt normal gene function, causing genetic disorders (17) (e.g., hemophilia A (18), Alström syndrome (19) and neurofibromatosis (20)). MEIs also occur in somatic tissues and have been linked to the initiation and development of certain cancers (21–23).

Given the biological significance of MEIs in genetic analysis, an increasing number of tools have been developed to explore MEIs in whole-genome sequencing (WGS) data as compared to a reference genome. VariationHunter (24) and Hydra (25), which were originally developed for detecting general structural variations, were fist transferred to MEI detection. More specific tools, including TEA (26), Retroseq (27), TIF (28), Mobster (29), Tangram (30), TEMP (31), T-lex2 (32), ITIS (33), RelocaTE2 (34), STEAK (35), MELT (36), TranSurVeyor (37), ERVcaller (38), TEBreak (39), and AluMine (40), were later designed to increase the quality of MEI identification. In general, most tools have attempted to explore MEIs in WGS data based on the statistics of hallmarks (e.g., discordant read pairs, clipped reads, ME consensus sequences, target site duplication, and poly A/T). These tools filtered candidates in the MEI calling process using different strategies. The reported recall rate based on each benchmark dataset ranged from 71.9–85.6%, and the reported precision were 88.3– 88.9%. However, only a few tools directly return the genotype of non-reference MEs. This inspired the design of TypeTE (41), which was built upon the calls produced by MELT. Another issue is that these tools all utilize hard-coded strategies to manually determine thresholds of the parameters from prior knowledge during detecting, filtering, and genotyping the MEIs. The recently published xTea first introduced a machine learning algorithm, Random Forest, into MEI genotyping with extracted features (42) and claimed improved recall rates of 88% for Alu, 93% for L1, and 86% for SVA. Clearly, significant improvements are still required to increase the accuracy of MEI calling and genotyping.

Deep learning algorithms hold great promise for tackling this issue. As widely adopted in the biomedical field in recent years (43), deep learning-based tools have improved performance in many bioinformatic methods, such as predicting splice junctions (44), enhancing Hi-C resolution (45), and determining scaffolding protein functional sites (46). Compared to other machine learning algorithms, deep learning algorithms can automatically capture elusive features and perform better when fed with sufficient training data. This advantage has been demonstrated by DeepVariant in detecting and genotyping of single nucleotide variants and small indels in WGS data (47). DeepVariant converted aligned reads to pileup images around candidate loci and trained an inception convolutional neural network (CNN) model to predict the probabilities of variant genotypes. It reported an F1 score of 99.98%, which outperformed the dominantly used GATK HaplotypeCaller (99.91%).

In the present study, we designed DeepMEI, a deep learning-based model for MEI identification and genotyping based on WGS data. DeepMEI uses soft-clipped and discordant reads to explore candidate insertion loci and then applies a trained a CNN model to predict the probability of non-reference ME genotypes. We benchmarked DeepMEI and existing tools against the datasets validated by Pacbio HiFi sequencing or PCR experiments, and that DeepMEI significantly improved recall rates and genotype accuracy. Finally, using DeepMEI, we uncovered abundant novel non-reference MEs in 3,202 high-coverage WGS samples from the phase 4 release of the 1000 Genome Project (1kGP), which can be explored for new insights in population genetics and biological studies of MEI.

## MATERIAL AND METHODS

### DeepMEI pipeline implementation

#### Identification of candidate non-reference ME loci

Possible MEIs were first identified across the sequencing data according to the consensus sequence of L1, Alu, and SVA. For each BAM or CRAM file, reads tagged with soft-clipping were extracted, and soft-clipped sequences were aligned to a library of ME consensus sequences provided by the Repbase (48). For soft-clipped sequences that did not meet the minimum length (<10 bp), the paired reads were substituted during the alignment. Positively matched reads were tracked for their genomic positions. Two or more ME-containing reads were grouped into a cluster when the distance between any two soft-clipped positions was shorter than 50 bp. Such clusters were considered candidate MEI loci, and, for each cluster, the median of the soft-clipping positions was considered as the insertion point. To reduce false positives due to alignment errors, candidate MEI loci were excluded if poly(N) sequences (>10 bp) were near the insertion points (within 30 bp).

#### Encoding aligned reads into pileup images

For each candidate MEI, reads that fully or partially aligned within 50 bp upstream or downstream of the insertion position were extracted from the BAM or CRAM file. The reads were divided into three groups: 1) no soft-clipping or > 5 bp distance between clipping position and insertion position; 2) soft-clipping occurred on the 5’ end of the read and the clipping position was within 5 bp upstream or downstream of the insertion position; and 3) soft-clipping occurred on the 3’ end and the clipping position within 5 bp upstream or downstream of the insertion position. Next, the grouped reads were sorted separately. In group 1 and group 3, reads were sorted in ascending order according to the start position recorded in the POS field of the BAM file. In group 2, reads were sorted in ascending order according to the end position that was calculated based on the POS and CIGAR of the BAM file.

After sorting, the read information was converted to a six-channel, 351 × 100 pileup image. Each pixel represented a base, and each channel represented a feature with the original value converted to grayscale, including nucleotide base, base quality, mapping quality, read strand, base matches reference sequence, and base matches ME consensus sequence. In the pileup image, the stack order was from the top to bottom as five repeats of the reference sequence, read group 1, read group 2, and read group 3. Random downsampling of the reads was performed if the image height exceeded 100.

#### Training of MEI genotyping model

The training dataset was constructed using ten samples of WGS data from the 1kGP phase 4 release, including HG00096, HG00097, HG00099, HG00100, HG00101, HG00102, HG00103, HG00105, NA12878, and NA12718 (49). Annotated structural variation calls were yielded from the VCF (freeze_V3) released by the project. According to the annotation information, non-reference MEs (Alu, L1, and SVA) were retrieved with a minimal insertion length of 50 bp. After manual confirmation for the presence and genotype using the Integrative Genomics Viewer (IGV) (50), valid MEIs were included in the positive dataset, while corresponding genomic positions without insertion in the rest of the samples were included in the negative dataset. Pileup images were generated as described previously for the training data.

The MEI classifier was built based on the Inception v3 architecture, a 42-layer CNN model that used RMSprop optimizer, label smoothing, factorized 7 × 7 convolutions, and an auxiliary classifier to improve the accuracy (51). Specifically, TensorFlow 2.0 was used to implement the Inception v3 architecture (52). The input layer was modified to receive the pileup image of size 351 × 100*6, and the model emitted probabilities for three genotypes of homozygous reference, heterozygous, and homozygous insertions. The highest probability determined the genotype of each MEI site. With a starting learning rate of 0.004, the model automatically reduced the learning rate until the val_loss stopped improving. The training was performed on a deep learning server with 1 TB memory, a 96-core CPU, and ten Nvidia Tesla V100 cards (Nvidia, Santa Clara, CA, USA) for 96 wall-clock hours. The model with the highest accuracy among the training process was selected as the final model.

#### Identification of MEI using DeepMEI

Sequencing data was aligned to the human reference genome using BWA (53). Candidate MEIs were screened as described. For each candidate MEI, a total of 41 pileup images were generated with the insertion point as centric and with stepwise shifting for ±20 bp of the insertion point. The results of DeepMEI were returned in a VCF format and included insert positions, length of TSD, breakpoints, poly A, family of ME, genotypes, and Phred quality scores logarithmically converted from genotype probabilities.

#### Using known MEIs to perform joint analysis

MEIs from the 1kGP phase 4 release using DeepMEI were converted into a bed file. The start and end position of the bed file is the corresponding breakpoints of each MEI. A joint analysis option was inbred and the bed file from known MEI can be adding to candidate MEI for further analysis. Joint analysis increased runtime from 38 minutes to 59 minutes. The way to perform joint for analysis of a cohort of multiple sample is different. User can merge bed file (e.g. HG002.bed) in DeepMEI output directory of all sample into a single file and use – b to generate a joint calling MEIs.

#### Perform a joint analysis using known MEIs

The MEIs from the 1kGP phase 4 release were identified using DeepMEI and converted into a bed file, where the start and end positions corresponded to the breakpoints of each MEI. The joint analysis option was then applied, and the bed file containing known MEIs was added to the candidate MEIs for further analysis. This joint analysis increased the runtime from 38 minutes to 59 minutes.Performing a joint analysis for a cohort of multiple samples requires a different approach. The user can merge the bed files (e.g., HG002.bed) from the DeepMEI output directory of all samples into a single file and use the “-b” option to generate a joint call result

### Generation of benchmark dataset and evaluation of DeepMEI performance

Two real-world benchmark datasets were constructed to evaluate DeepMEI and existing tools. The first benchmark dataset was created using high-confidence structural variation (SV) in HG002 called by the Genome in a Bottle (GIAB) Consortium using short-read and long-read sequencing techniques. From the VCF file (HG002_SVs_Tier1_v0.6.vcf.gz), non-reference L1, Alu, and SVA were extracted based on annotation of the insertion sequence by Repeatmasker (4.1.2). The existence and genotype of each MEI was confirmed using the PacBio HiFi sequencing BAM file (HG002.Sequel.10kb.pbmm2.hs37d5.whatshap.haplotag.RTG.10x.trio.bam) by IGV and Repeatmasker. The other benchmark dataset, confirmed by PCR-based assays in a study by Payer et al. (54), included 7,898 non-reference Alu insertions on 145 polymorphic sites in 90 CEU HapMap samples. The original MEI positions were converted from hg18 to GRCh38 using Liftover (55).

The performance of DeepMEI was compared to five commonly used tools, MELT (36), RetroSeq (27), ERVcaller (38), Mobster (29), and xTea (42). The HG002 BAM file HG002.hs37d5.60X.1.bam was downloaded from the GIAB FTP site, and, for benchmark dataset 2, the CRAM files were from ftp://ftp.sra.ebi.ac.uk/vol1/run/. Each tool was run with the respective parameters and ME reference sequences provided. Since RetroSeq generates excessive insertion calls, only those with FL = 6 & GQ ≥ 28, FL = 7 & GQ ≥ 20, or FL= 8 & GQ ≥ 20 were kept according to the filtering method of 1000-Genome-CEU-Trio-Analysis in the RetroSeq document. A MEI called within 50 bp of the benchmark position was considered positive. Precision, recall, and F1 statistics were used to evaluate the performance.

### Analysis of high-coverage 1kGP samples using DeepMEI

High coverage WGS CRAM files of 3202 samples from the 1kGP phase 4 release were downloaded from the project cloud server and analyzed using the DeepMEI pipeline. Called MEIs were annotated using the Ensembl Variant Effect Predictor (56). Frequencies of non-reference MEs were calculated, and population stratification analysis was performed using principal component analysis. All analyses were performed using R Statistical Software (v4.1.2) unless otherwise specified.

## RESULTS

### DeepMEI pipeline overview

To improve the accuracy of MEI identification, we developed DeepMEI, which utilized a deep CNN model to recognize non-reference MEs from WGS data. In brief, DeepMEI consists of candidate detection and genotyping modules (Figure 1). The detection module scans soft-clipped and discordant reads across entire BAM files for ME fragment sequences to obtain candidate MEI loci. Aligned reads around each candidate MEI are grouped, sorted, and encoded into a pileup image. Next, the genotyping module predicts the genotypes of candidate MEIs by a trained CNN model based on the Inception V3 architecture, which can automatically extract image features and return accurate classification similar to manual visualization. The DeepMEI pipeline takes an aligned BAM file as an input and produces the genomic position, ME type, genotype, and Phred quality score of MEIs in the VCF format as an output.

**Figure 1.**
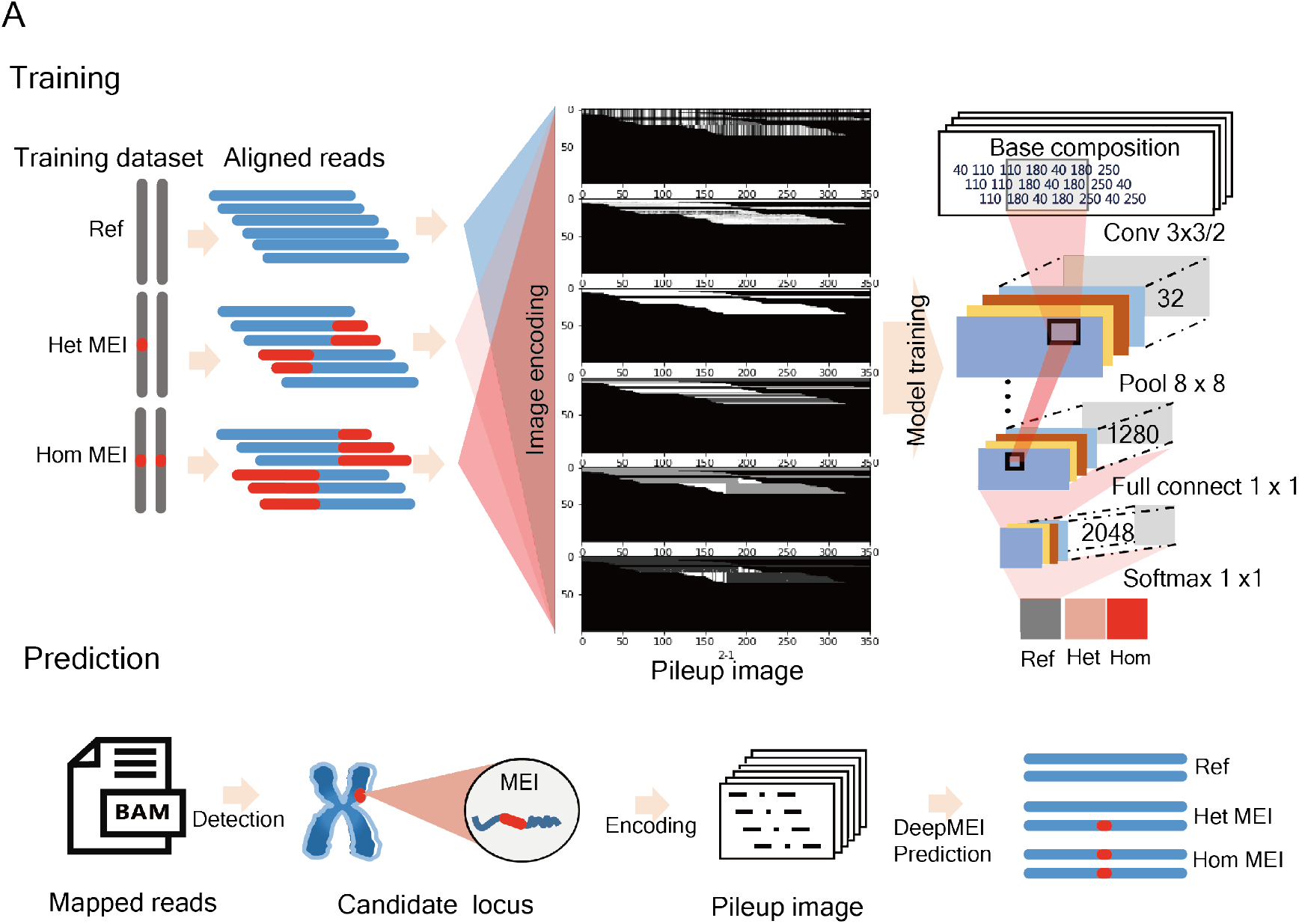
Overview of DeepMEI workflow. **A)** DeepMEI converted aligned reads into a pileup image around the non-reference ME sites and trained a convolutional neural network model to identify MEIs and call the genotypes. **B)** To call MEIs from given samples, DeepMEI first identified candidate insertion sites from short-read sequencing data. Next, DeepMEI converted aligned reads around the candidate sites into pileup images. Finally, the trained CNN model predicted the probability of candidate MEI genotypes and returns the results.

### Benchmarking DeepMEI and existing tools

We generated a highly credible benchmark dataset using HG002 released by GIAB. SV calls of HG002 were generated by integrating high coverage, short- and long-read genome sequencing data and multiple SV calling tools. Alu, L1, and SVA in the HG002 SV call set were labeled using Repeatmasker. The existence and genotype of each ME was further confirmed by checking PacBio HiFi sequencing data with IGV and Repeatmasker. The final dataset contained 2320 non-reference MEIs across all nuclear chromosomes, including 1,818 Alu, 392 L1, and 110 SVA. Among all Alu insertions, AluYa5, AluYb8, and AluY were predominant (68.4%, Figure S1), which were reported as active MEs (57). The rates of heterozygous insertion were 66–79% in the youngest AluY lineage, higher than the rates in the ancient AluJ and AluS elements (36–48%). Similar correlations were also observed in L1 and SVA (Figure S1).

Performance of DeepMEI was compared to MELT (v2.2.2), Mobster, Retroseq, ERVcaller (v1.4), and xTea. Of note, Mobster and Retroseq only provide calling sites without genotypes. In order to perform a fair comparison of the tools, all tools with the most recent version were used for two evaluations based on truly called MEI sites with or without considering genotype. Figure 2A illustrates the precision, recall, and F1 score calculated based on truly called sites without considering genotype.

**Figure 2.**
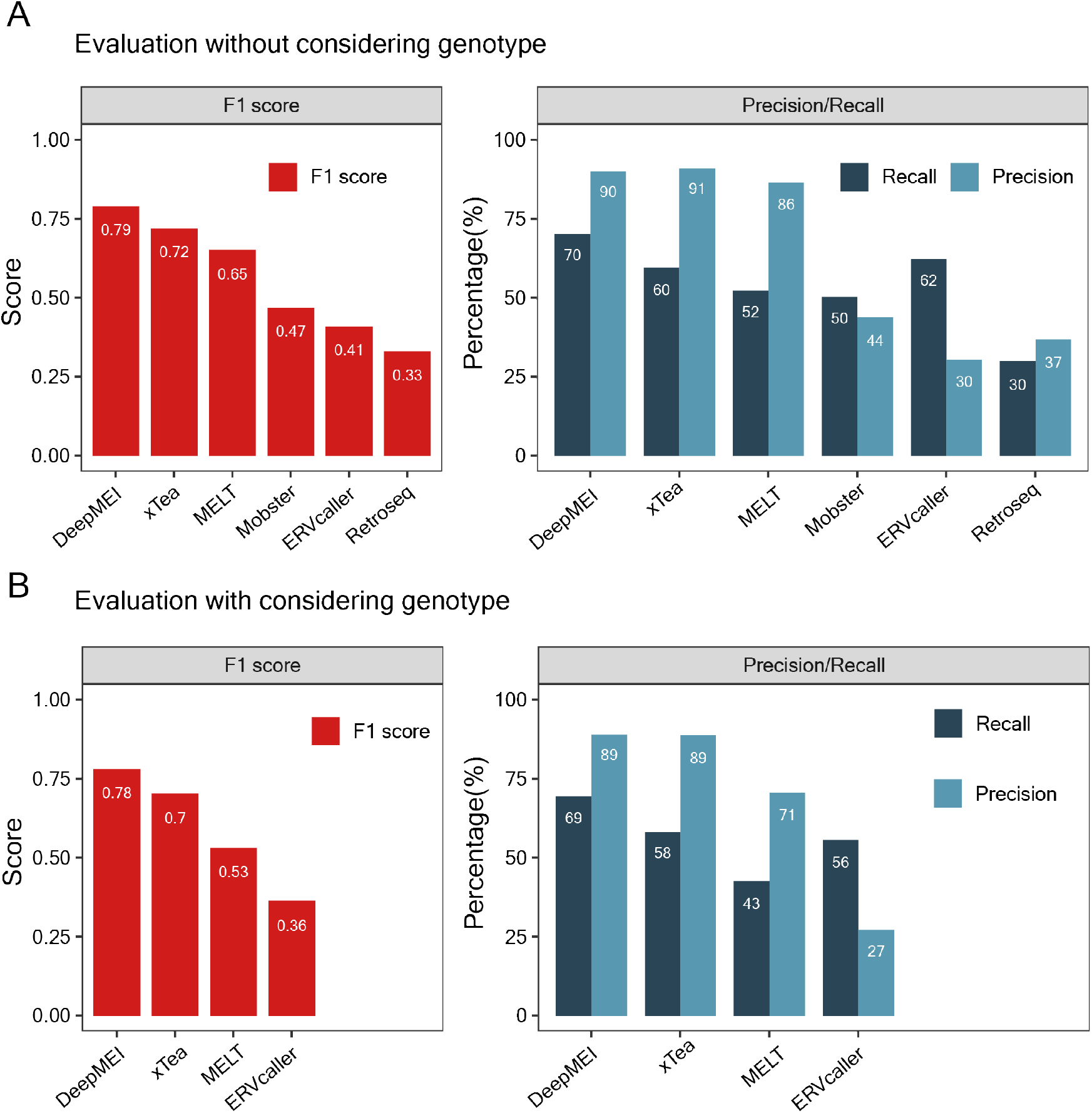
Comparison of DeepMEI performance with five existing tools for analyzing the GIAB HG002 benchmark dataset. **A)** the first evaluation was based on the called MEI sites without considering genotype information. **B)** the second evaluation was based on the called MEI sites considering correct genotype information. Genotype output was unavailable in Mobster and Retroseq; therefore, the two tools were not included in the genotype-based evaluation.

DeepMEI outperformed other tools in terms of the overall performance with an F1 score of 0.789, which was followed by xTea with an F1 score of 0.72. The precision rate of DeepMEI (90.1%) was 1% lower than xTea (91%), but DeepMEI discovered significantly more MEIs with a recall rate of 70.3% compared to xTea’s 59.6%. Following DeepMEI and xTea, MELT yielded a precision of 86.5% and a recall of 52.3%. ERVcaller had the highest recall of 62.3% among all tools, whereas the precision rate of 30.4% was unsatisfactory. When evaluating the truly called MEI sites with correct genotype information, DeepMEI and xTea remained the top performing callers; of them, the recall and precision rates were slightly decreased for 1–2% (Figure 2B). MELT had a significant reduction in recall from 52.3% to 42.7% and in precision from 86.5% to 70.6%, as it is known to be effective in capturing non-reference MEs but performs poorly in genotyping (41). Moreover, we compared the tools for Alu, L1, and SVA separately, and DeepMEI was still found to perform better than other tools (Figure S2–S4).

In the recalled non-reference MEIs, the genotyping errors of DeepMEI (1.1%) and xTea (2.4%) were significantly lower than those of ERVcaller (14%) and MELT (18.9%; Figure 3A). The genotyping errors included two types: called homozygotes on true heterozygous MEIs and called heterozygotes on homozygous MEIs. Both types of errors were found in DeepMEI and xTea. The majority (88.9%) of DeepMEI’s genotyping errors were heterozygous MEIs wrongly called as homozygotes, while xTea was the opposite—70.8% of xTea’s errors were homozygotes wrongly called as heterozygotes. ERVcaller and MELT also showed abundant errors in genotyping homozygous MEIs.

**Figure 3.**
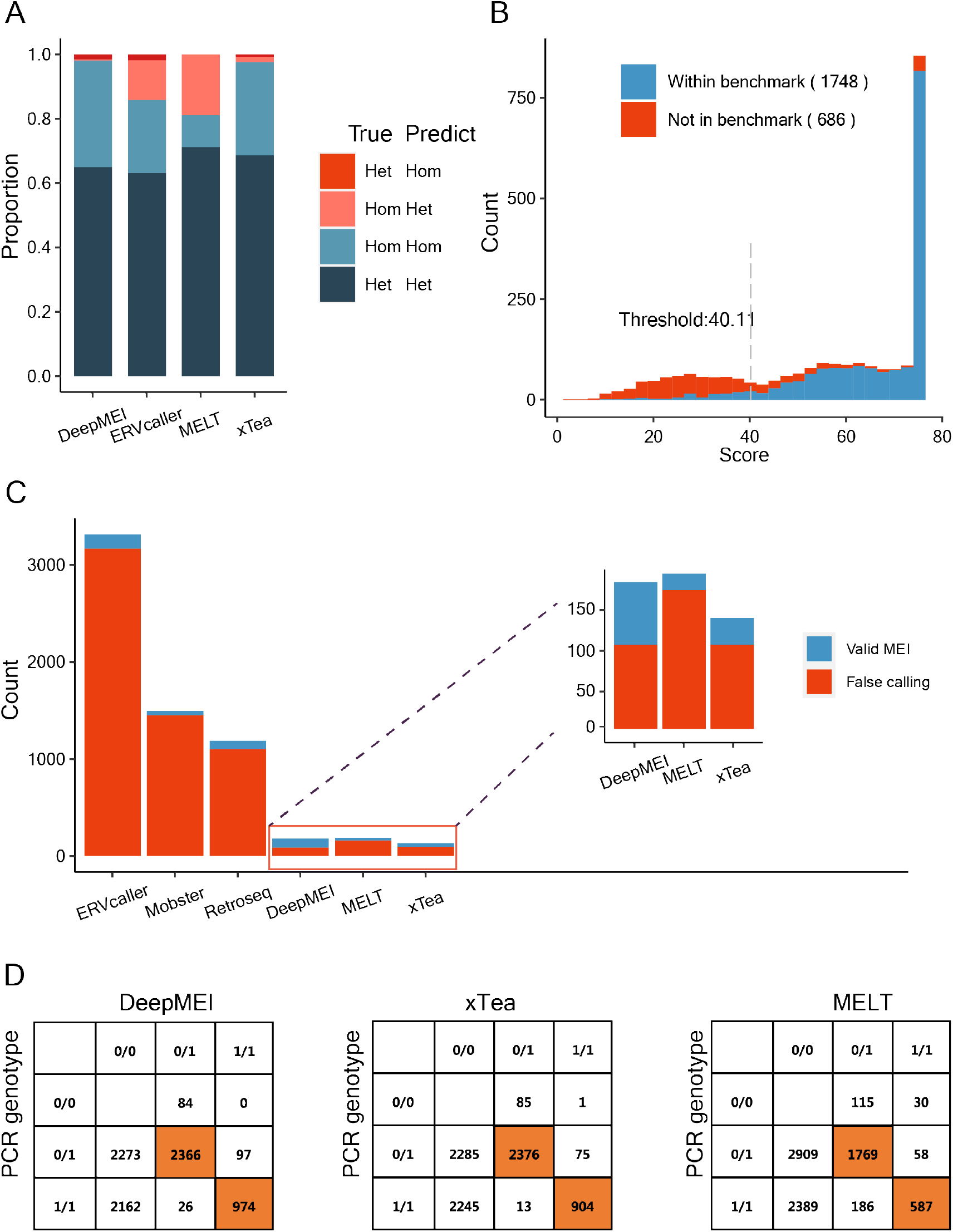
Performance comparison of DeepMEI with MELT, xTea, and ERVcaller on genotyping accuracy. **A)** DeepMEI and xTea showed low genotyping error rates in recalled non-reference MEs of the GIAB HG002 benchmark dataset. **B)** DeepMEI showed the Phred quality scores for all candidate sites. A score > 40 was set as the cutoff for quality filtering. **C)** DeepMEI and existing tools identified true non-reference MEs that were not in the GIAB HG002 benchmark data release. **D)** Performance comparison of DeepMEI with xTea and MELT on a PCR-based benchmark dataset from a study by Payer et al(54).

DeepMEI output a Phred quality score for each MEI call to indicate the confidence level. In the benchmark dataset, 73.1% of MEIs yielded a Phred quality score > 40 (Figure 3B). However, of the 187 “false positive” MEIs, which were called by DeepMEI but not in the benchmark set, 1810 had a Phred quality score > 40. We further assessed the “false positive” MEIs using PacBio HiFi sequencing data of HG002 released by GIAB. In the 187 “false positive” MEIs called by DeepMEI, 77 (42.5%) were de facto true calls missed by the benchmark dataset (Figure 3C). Correcting such calls could increase the precision rate of DeepMEI to 94.3%. xTea also identified 33 (27%) true MEIs among “false positive” calls, and the corrected precision rate was 94.1%. For MELT, Mobster, ERVcaller, and Retroseq, 170–3195 “false positive” MEIs were recorded, and only 20–121 (2.5–10.5%) were true insertions, which had a mild effect on the precision rates of the four tools.

We next compared the performance of DeepMEI, xTea, and MELT using another benchmark dataset reported in a study by Payer et al. (54). This dataset consisted of 175 high-frequency Alu sites in 90 samples from 1kGP. All 4,736 heterozygous and 3,162 homozygous insertions were experimentally validated using a PCR-based assay (Figure 2C). By screening the corresponding sites in CRAM files of the 1kGP phase 4 release, DeepMEI identified 3,463 (43.8%) non-reference Alu insertions, of which 96.4% were genotyped correctly. Similarly, xTea discovered 3,368 (42.6%) Alu insertions, and 97.4% were correctly genotyped. Comparing the results, 3,157 (88.7%) insertions were consistently called by DeepMEI and xTea, whereas 4,340 (60%) insertions on 109 polymorphic sites were missed by both tools. MELT showed fewer false positive calls (n = 2,600) and correctly genotyped 2,356 (91%) insertions.

### Tool concordance and performance in combination

Creating ensembles of multiple tools is a common strategy in calling structural variation. It is thought that incorporating the results of several tools can improve the sensitivity or specificity over using only a single tool. We investigated the concordance of the tools and the performance of tool combinations (Figure 4). Among all 2,320 non-reference MEIs in the HG002 dataset, 1,882 were called by at least one tool, while only 490 were called by all tools. Intriguingly, 438 MEIs were not called by all tools, suggesting it is difficult to detect such MEs in short-read sequencing data. The concordance between any two tools ranged from 10.4–66.2% (Figure S9). In calling a total of 2,004 MEs, DeepMEI and xTea had 66% overlap, of which 1,720 were true MEs and 284 were false positives (Figure 4C). We compared all possible tool ensembles by requiring a call to be made by at least one caller. “DeepMEI+xTea” performed best with an F1 score of 0.81 but was very closely followed by using DeepMEI alone (Figure 4A). The other top tool ensembles were “DeepMEI+xTea+MELT”, “DeepMEI+MELT”, and “MELT+xTea”, all of which outperformed using xTea or MELT alone. Any tool ensemble involving ERVcaller, Mobster, or Retroseq performed poorly, consistent with the benchmarking results of these tools.

**Figure 4.**
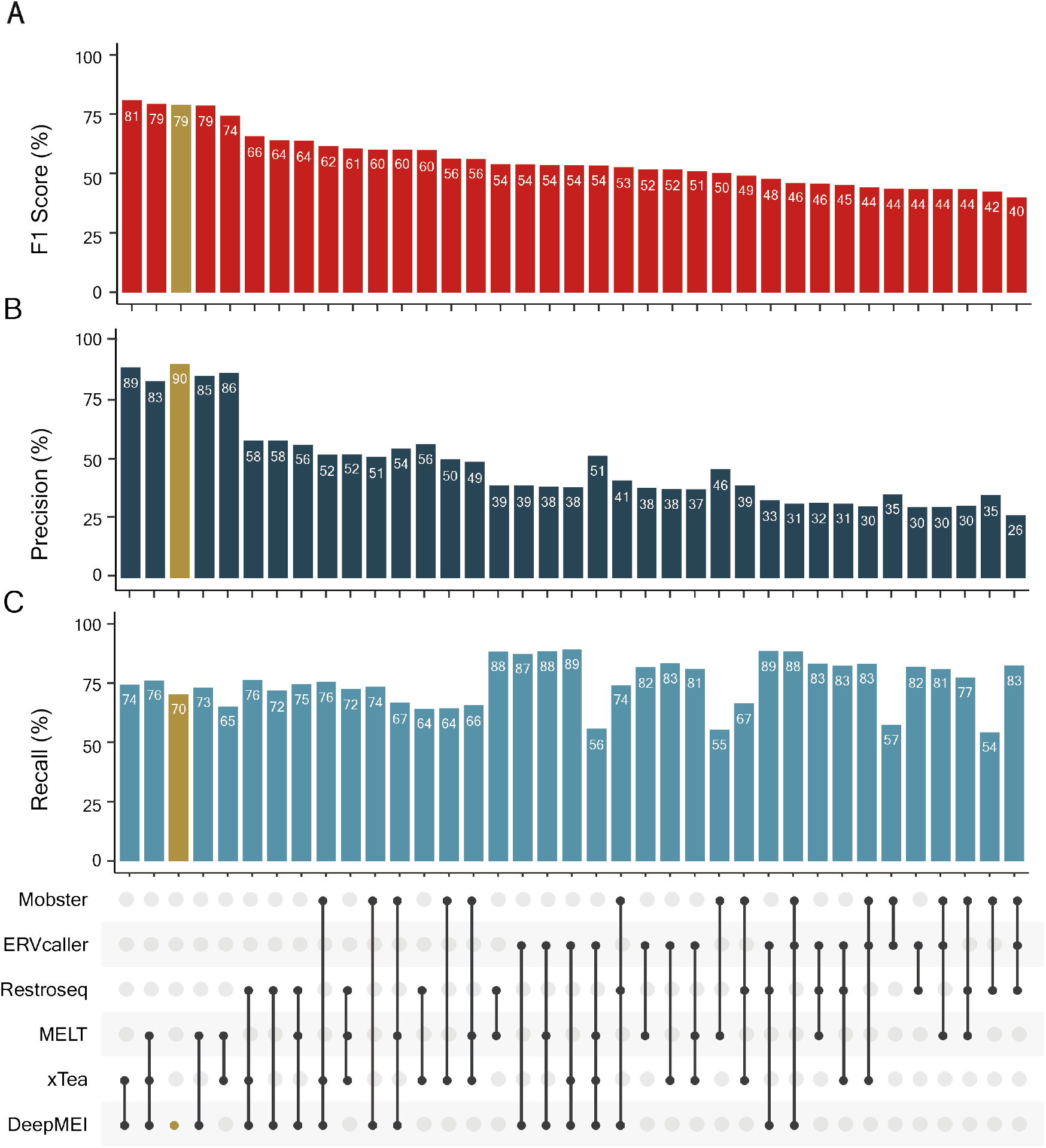
Comparison of DeepMEI’s performance with combinations of tools on the GIAB HG002 benchmark dataset. The top three combinations all included DeepMEI and performed similarly to DeepMEI alone.

### Depth, mappability, and ME integrity effect on performance

Earlier studies indicated that the sequencing platforms and genomic context can influence the calling of non-reference MEs (42). Here, we assessed the effects of sequencing depth, genome mappability, and ME integrity on performance. Using random down-sampling on the aligned reads of HG002, a set of BAM files with varied mean sequencing depth was generated and analyzed using DeepMEI, MELT, and xTea. As shown in Figure 5A–C, when the sequencing depth rose from 20X to 35X, the F1 score and recall rate of all three tools increased rapidly, while the precision rate remained almost unchanged. The increase in the recall rate slowed down after 35X, indicating that this depth Was the minimum for detecting MEIs in WGS data. DeepMEI had higher recall rates and F1 scores than xTea and MELT at all depths. DeepMEI also outperformed MELT and xTea on each ME type (Alu, L1, and SVA) at different depths, as shown in Figure S5.

**Figure 5.**
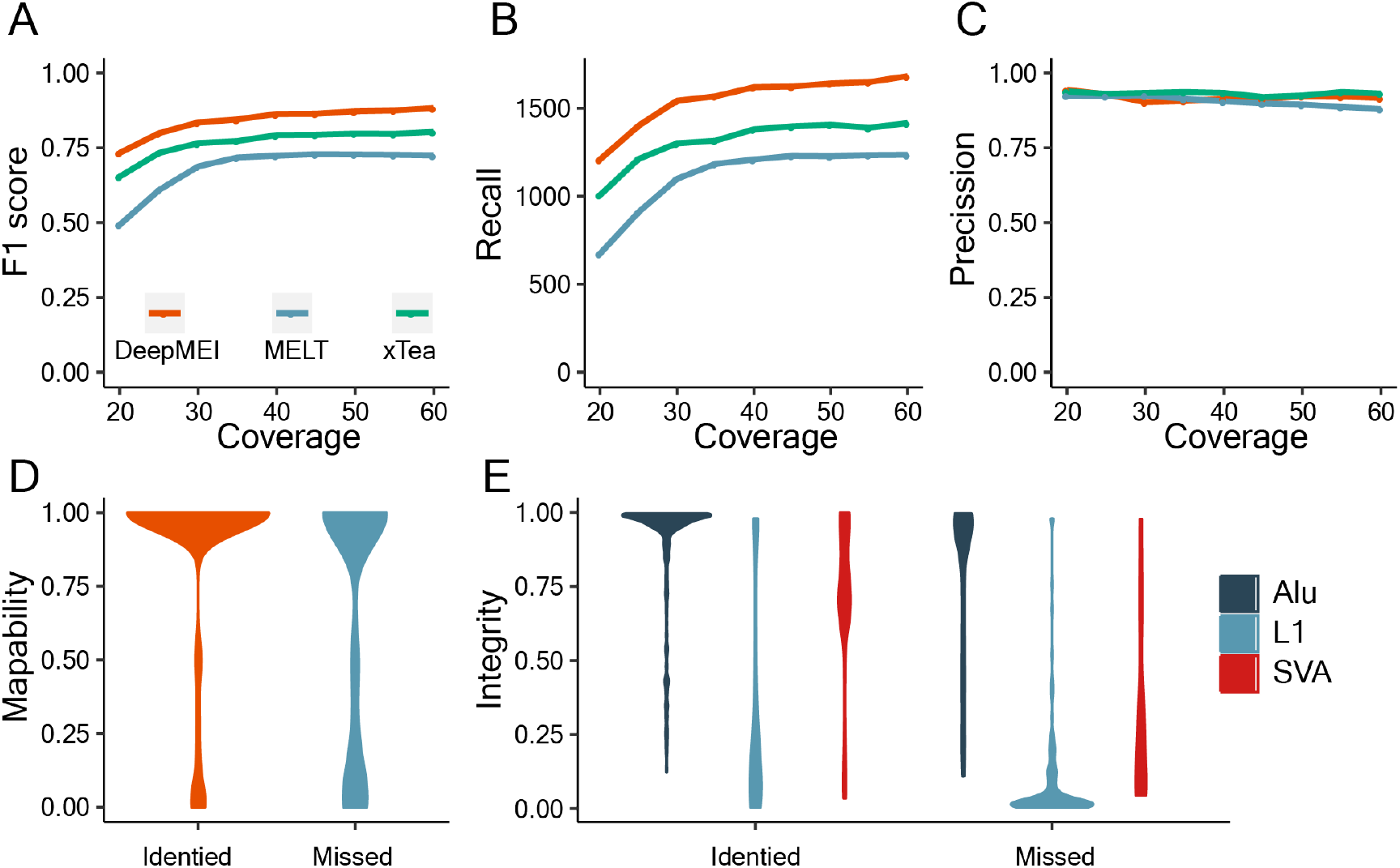
Evaluation of the factors affecting MEI identification. **A)** Increasing sequencing coverage improved overall performance of DeepMEI, xTea, and MELT. **B)** Coverage affected the recall rate of DeepMEI, xTea, and MELT. **C)** The influence of coverage on precision rate was limited. **D)** The performance of DeepMEI was reduced in genomic regions with low mappability. **E)** ME integrity correlated with the recall rate of DeepMEI.

A highly homological and polymorphic sequence reduces genome mappability for short-read sequencing, which is crucial for the interpretation of WGS data. The genome mappability score (GMS), as a measure of genome complexity, is a weighted probability that any read could be unambiguously mapped to a given position (58). We assessed the correlation between GMS and DeepMEI’s performance (Figure 5D). In the HG002 dataset, 78.4% (1,819) of non-reference MEs had a mean GMS > 0.75 on the 300 flanking bases, suggesting a high uniqueness, and 21.3% (493) of MEs had a GMS < 0.25, indicating regions of low mappability. In the HG002 dataset, DeepMEI identified 1397 (76.8%) of the MEs with a GMS > 0.75, and only 50.1% (247) of the MEs with a GMS < 0.75 were called by DeepMEI. This demonstrated that the calling capability of DeepMEI significantly correlated with mappability (Fisher’s exact P < 5.311e^-7^). Similar results were seen in xTea and MELT (Figure S5). It is notable that 18.3–31% of the benchmarking MEs with GMS of 1.0 were missed by all three tools.

Earlier studies reported that a significant portion of non-reference MEs in the human genome are truncated, indicating deactivated retrotransposition ability (3, 59). The integrity of MEs implies that they contain information important in evolution, but truncated MEs may be neglected by callers. Using long-read sequencing data, we calculated integrity scores for each benchmarking ME as the percentage of the consensus sequence that was covered by a given inserted ME. In the HG002 benchmark dataset, most of the Alu elements (80.2%) were relatively intact (integrity score > 0.8). In the DeepMEI results, 84.8% and 66.6% of identified and missed Alu elements had an integrity score > 0.8 (Figure 5E). In contrast, 73.2% of the L1s were largely truncated (integrity score < 0.2), which was consistent with previous studies (42). DeepMEI identified 81.9% of the L1s with an integrity score > 0.2 and missed 59.2% of the L1s with an integrity score < 0.2. SVA elements showed the strongest effects of integrity on performance. In the DeepMEI results, 88.6% of identified SVAs had an integrity score > 0.2, and 75% of missed SVAs had an integrity score < 0.2.

### Identification of MEIs in 1kGP phase 4 release data

The recent release of 1kGP phase 4 data included high-coverage WGS (30X) for 3202 samples (49). Along with the sequencing data, 1kGP also generated a call set using GATK-SV, an ensemble caller refining MEI calling based on the output of MELT. We analyzed the 1kGP phase 4 data using DeepMEI, and the MEI calls were compared with the call set released by 1kGP. A total of 6,218,088 non-reference MEs were detected by DeepMEI in 3,202 samples, including 4,982,745 Alu, 943,427 L1, and 291,916 SVA insertions, which was about 1.71-fold more than the 1kGP MEI call set (3,334,832 total MEs; 2,972,136 Alu; 254,528 L1; 108,168 SVA). DeepMEI recalled 85.7% of Alu, 78.5% of L1, and 82.8% of SVA insertions. Notably, the DeepMEI call set corrected the genotypes for 10.4% of MEI sites that overlapped with 1kGP, showing a significant improvement in genotype accuracy as demonstrated in previous evaluations. For each sample, a mean of 1,558 Alu, 291 L1, and 93 SVA insertions were found by DeepMEI (Figure 6A and Figure S6). African populations (AFR) harbored more MEIs (2233.2 per sample) than East Asian (EAS; 1847.5 per sample), South Asian (SAS; 1871.4 per sample), European (EUR; 1829.8 per sample), and American populations (AMR; 1877.0 per sample). DeepMEI calling results of the 1kG phase 4 data can be publicly accessed as described in the Material and Methods section.

**Figure 6.**
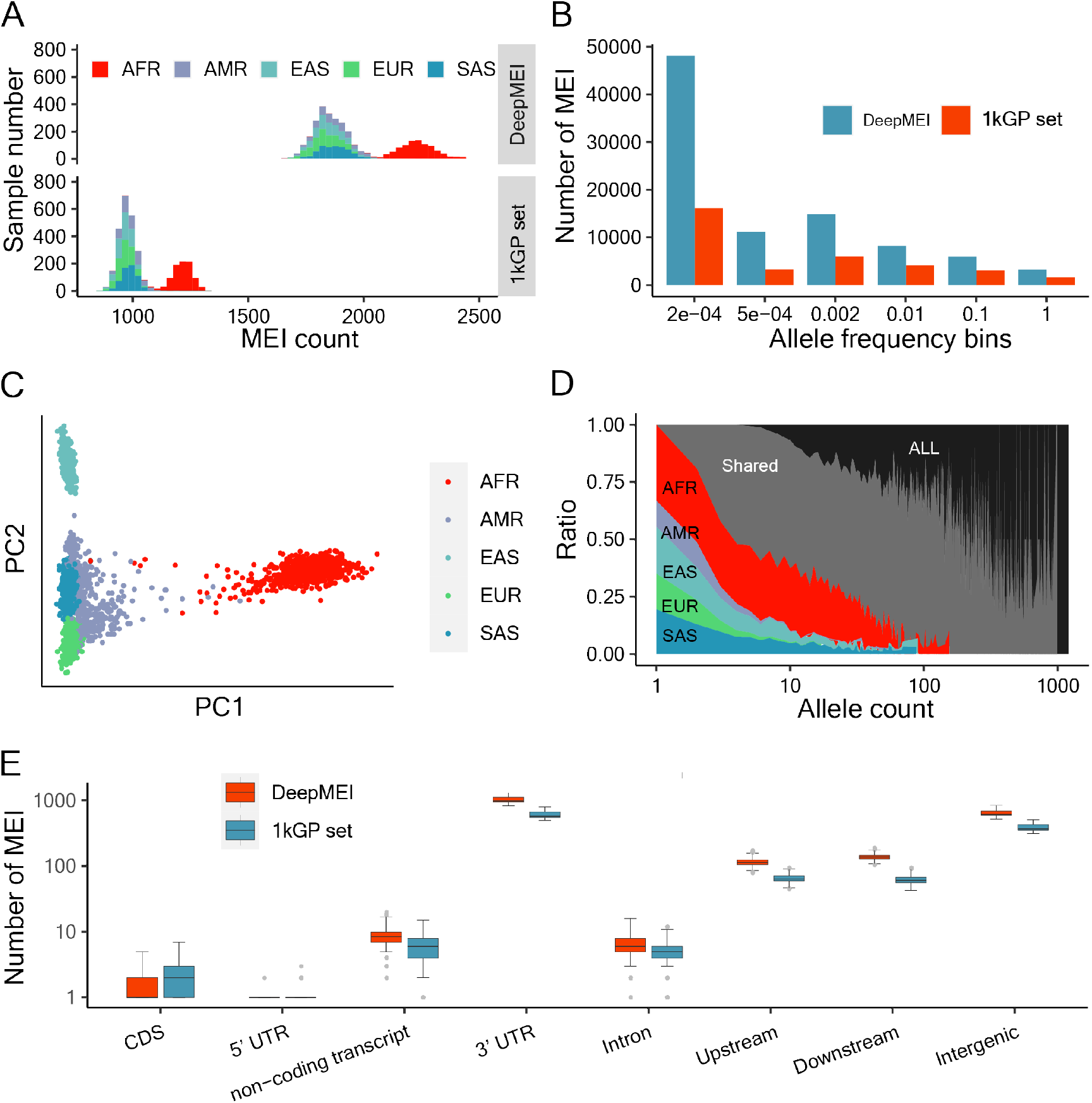
Reanalysis with DeepMEI of the 1000 Genome Project (1kGP) phase 4 release data. **A)** DeepMEI identified more non-reference MEs in the 1kGP samples than the original release. African samples showed more non-reference MEs than samples from other populations. Population abbreviations are AFR: Africa, AMR: America, EAS: East Asia, EUR: Europe, SAS: South Asia. **B)** Allele frequency distribution of DeepMEI and 1kGP call sets; **C)** Population stratification analysis using Alu insertions as markers stratified 1kGP continental populations. **D)** Unique and shared non-reference MEs identified in different populations; the AFR data harbored more unique non-reference MEs than data from other populations. **E)** Enrichment analysis showed depletion of non-reference MEs in coding sequences, 5’ UTR, and 3’ UTR regions. **F)** Distributions of non-reference MEs count in genome regions

Next, we calculated the allele frequency of non-reference MEIs in the 1kGP cohort using 2504 unrelated samples (Figure 6B). Most (89.9%) MEIs were rare (allele frequency < 0.01), among which 43,627 were singletons and 13,607 were doubletons. We performed population stratification analysis using Alu insertions. The results showed that Alu insertions were able to clearly differentiate between all populations (Figure 6C). Notably, AFR samples harbored significantly more unique non-reference MEs than non-AFR samples (Figure 6D and Figure S7). AFR-specific MEIs accounted for more than 13.4% of all MEI insertions until the allele count reached 48. This was consistent with previous reported higher MEI diversity in AFR samples compared with other populations (60). With an increase in allele count, the shared MEI insertions among populations gradually increased, and population-specific MEIs showed a decreasing trend.

We assessed the functional effects of MEIs in the 1kGP cohort. Consistent with earlier studies (15), the majority (86%) of the non-reference MEIs were in intronic and intergenic regions (Figure 6E, Figure S8), suggesting that MEIs in these regions are more likely to be fixed in the genome.

## DISCUSSION

In the present study, we presented DeepMEI, a deep CNN model-based tool to identify non-reference MEIs using WGS data. Different from existing tools that detect MEIs by setting thresholds on multiple features derived from candidate MEIs, DeepMEI followed an entirely different strategy that mimics how humans recognizes MEIs and made genotype calling decisions according to IGV screenshots. DeepMEI transformed the aim of MEI identification into image recognition, as inspired by DeepVariant (47), and outperformed the five existing tools MELT, RetroSeq, ERVcaller, Mobster, and xTea on a high-quality benchmark of HG002 data generated from GIAB. Among all of the tools, DeepMEI achieved the best overall performance (F1 score of 0.79), precision rate (0.90), and recall rate (0.7) in MEI identification. We also demonstrated that DeepMEI had higher sensitivity and comparable precision in genotyping MEIs than xTea, and both DeepMEI and xTea outperformed MELT, which has been widely used in large genome studies (49, 60, 61). Using DeepMEI, we reanalyzed MEIs in 3,202 high-coverage WGS samples from the phase 4 data release from 1kGP data and uncovered 836– 1419 more non-reference MEIs per sample compared to the original release (49), demonstrating the effectiveness of DeepMEI and providing a more comprehensive MEI resource to study their roles in human disease.

In WGS data, non-reference MEIs demonstrate several features that were utilized here for identification. ME consensus sequences, polyA/T signals, and target site duplications can be observed on the clipped or discordant reads around the insertion site. Most existing tools capture these features and optimize a set of manually determined parameters to call candidate MEIs and filter for valid ones. However, designating a unitary threshold value (e.g., the count of reads supporting the presence of an MEI) across the genome is difficult to apply to variable real sequencing data, which would in turn hamper recall and precision. Thereby, recently developed tools, such as xTea and gnomAD-SV, have applied Random Forest models in genotyping modules to assemble features in the context of local sequencing and genome information, which significantly improved their accuracy (42, 61). DeepMEI also explored candidate MEIs across the genome using sequence features of clipped and discordant reads to reduce the computational cost. However, different from existing tools, DeepMEI converts aligned reads around the candidate loci to a pileup image and transforms genotyping question into image recognition. Using the Inception v3 architecture, DeepMEI optimized under 100 MB of parameters implemented in this 42-layer pre-trained CNN model to automatically extract features from the pileup image, thereby avoiding the issue of subjective inference in parameter and threshold selection. DeepMEI was designed to tolerate loose criteria for candidate MEI inclusion, which significantly improved recall, and to use a robust deep learning model to accurately determine the likelihood of the existence and genotype of MEIs. Therefore, DeepMEI identified many true new MEIs that were not detected by any of the other tools. Nevertheless, about 18.9% of the MEIs validated by long-read genome sequencing or PCR assays were missed by DeepMEI and the other existing tools. Part of these MEIs were not identified during MEI candidate detection, as ME consensus sequences were absent on the soft-clipped reads around the insertions. Other failures were due to low-complexity genome sequence near the insertions, which can cause misalignment of the reads to other ME loci in the genome and lead to insufficient reads supporting the presence of the MEI. The “missing” MEIs will guide us to fine-tune the model in future updates.

Practically, previous studies often combined several tools to increase sensitivity. We assessed the performance of tool combinations and found that the top performer was the combination of DeepMEI + xTea, with a recall of 0.74, which was 4% higher than using DeepMEI alone. However, the precision also declined by 1%. As a result, the overall performance of “DeepMEI + xTea” was only 2% higher in F1 scores than using DeepMEI alone. Moreover, DeepMEI was in the top four combinations with F1 scores ranging from 0.79 to 0.81. The precision dropped when adding more tools, but the improvement in recall was small. We also assessed other factors that may affect MEI identification. Interestingly, a high mean coverage had a mild negative influence on the precision of DeepMEI, xTea, and MELT, whereas a low coverage depth (<30X) greatly reduced the recall (Figure 5B). That is, most of the MEIs obtained by MELT from the low-depth WGS data from 1kGP are highly credible, except for their genotypes (15). From the present evaluation, 35X to 40X coverage was found to be sufficient for MEI identification. Another factor that affected DeepMEI’s performance was genome mappability (Figure 5D). Genome mappability measures the uniqueness of each region in the reference genome. Low genome mappability relates to highly repetitive or polymorphic sequences. Since genome mappability mostly depends on the length of the reads produced by the sequencing experiment, improving the sensitivity of MEI detection in regions with low mappability will rely on long-read genome sequencing.

The recent phase 4 data release from 1kGP published sequencing and variant data of 3202 WGS samples with a coverage ranging from 27X to 71X, providing a valuable resource for understanding the genomic diversity of 26 populations across the world. In the original dataset, MEIs in these samples were discovered using the MELT algorithm. The presence of MEIs may be underestimated given the relatively low sensitivity of MELT, as demonstrated in the present evaluation and earlier studies (42). Here, DeepMEI called 4,982,745 Alu, 943,427 L1, and 291,916 SVA insertions across the 3,202 samples. Compared to the original release, a cohort-level increase of 2,883,256 MEIs was reflected in our re-called dataset, and an average of 900 MEIs per genome were increasingly identified. Up to 92.2% of the increment was from rare MEIs with a MAF < 0.01, which could advance future studies for understanding the impact of rare MEIs respect to human disease or phenotype and evolution. Population stratification analysis based on the DeepMEI call set yielded similar separation effects as using single nucleotide variants. African populations exhibited more diverse and unique MEIs than other populations, consisted with previous reports (60).

In conclusion, DeepMEI utilizes a novel image recognition strategy in MEI identification and offers significant improvements of recall and genotyping accuracy over existing tools. Notably, DeepMEI showed enhanced ability for finding rare MEIs, which have been presumably ignored in previous population genomics studies. We regenerated the MEI call set for the phase 4 data release from 1kGP high-coverage WGS samples and anticipate that this resource will benefit the community to further understand the functional and regulatory influences and the evolution of MEIs in humans. For future updates, we intend to make DeepMEI compatible with long-read sequencing data, to support other mobile element families, and to improve applicability to genotype somatic or polyploid changes in all tissues and species.

## DATA AVAILABILITY

The DeepMEI and datasets utilized in this study can be obtained from the following URLs.

DeepMEI source code: https://github.com/xuxif/DeepMEI

Training dataset: https://github.com/xuxif/DeepMEI/tree/main/DeepMEI/training_dataset

Benchmark dataset 1: https://github.com/xuxif/DeepMEI/tree/main/DeepMEI/benchmark_dataset

Benchmark dataset 2:

https://www.pnas.org/doi/suppl/10.1073/pnas.1704117114/suppl_file/pnas.1704117114.sd04.xlsx

GIAB HG002 sequencing and calling data: ftp://ftp-trace.ncbi.nlm.nih.gov/giab/ftp/data/AshkenazimTrio/analysis/NIST_SVs_Integration_v0.6/HG002_SVs_Tier1_v0.6.vcf.gz, ftp://ftp-trace.ncbi.nlm.nih.gov/giab/ftp/data/AshkenazimTrio/HG002_NA24385_son/NIST_HiSeq_H002_Homogeneity10953946/NHGRI_Illumina300X_AJtrio_novoalign_bams/HG002.hs37d5.60X.1.bam, ftp://ftp-trace.ncbi.nlm.nih.gov/giab/ftp/data/AshkenazimTrio/HG002_NA24385_son/PacBio_CCS_10kb/alignment/HG002.Sequel.10kb.pbmm2.hs37d5.whatshap.haplotag.RTG.10x.trio.bam

1kGP phase 4 sequencing and calling data (including 10 samples for the training dataset and 90 CEU samples for benchmark dataset 2): http://ftp.1000genomes.ebi.ac.uk/vol1/ftp/data_collections/1000G_2504_high_coverage/working/20210124.SV_Illumina_Integration/1KGP_3202.gatksv_svtools_novelins.freeze_V3.wAF.vcf.gz, ftp://ftp.sra.ebi.ac.uk/vol1/run/

DeepMEI calling results of 1kGP phase 4 samples:

https://github.com/xuxif/DeepMEI/tree/main/DeepMEI/1000g_high_callset

## ACKNOWLEDGEMENT

We thank Bo Yang, Tao Zheng, and other colleagues in the Information Center of West China Hospital for helping us operate and maintain computing resources. We thank Feng Zhu, Shunmin He and Bo Li for valuable comments in the data analysis and critical review of the manuscript. We thank the 1000 Genomes Project (Phase 4), the Genome in a Bottle Consortium, and colleagues for providing sequencing data of their samples.

## AUTHOR CONTRIBUTIONS

X.X. and F.B. conceived the study. X.X. created and evaluated the model, with contributions by X.W., Y.H., and C.J. The manuscript was written by F.B., with contributions by X.X. Study supervision and funding acquisition were conducted by F.B. and H.Y.

## SUPPLEMENTARY DATA

Supplementary Data are available at NAR online.

## REFERENCES

1. Cordaux, R. and Batzer, M.A. (2009) The impact of retrotransposons on human genome evolution. Nat Rev Genet, 10, 691–703.

2. Lander, E.S., Linton, L.M., Birren, B., Nusbaum, C., Zody, M.C., Baldwin, J., Devon, K., Dewar, K., Doyle, M., FitzHugh, W., et al. (2001) Initial sequencing and analysis of the human genome. Nature, 409, 860–921.

3. Goodier, J.L. (2016) Restricting retrotransposons: a review. Mob DNA, 7, 16.

4. Hata, K. and Sakaki, Y. (1997) Identification of critical CpG sites for repression of L1 transcription by DNA methylation. Gene, 189, 227–234.

5. De Cecco, M., Criscione, S.W., Peckham, E.J., Hillenmeyer, S., Hamm, E.A., Manivannan, J., Peterson, A.L., Kreiling, J.A., Neretti, N. and Sedivy, J.M. (2013) Genomes of replicatively senescent cells undergo global epigenetic changes leading to gene silencing and activation of transposable elements. Aging Cell, 12, 247–256.

6. Newkirk, S.J., Lee, S., Grandi, F.C., Gaysinskaya, V., Rosser, J.M., Vanden Berg, N., Hogarth, C.A., Marchetto, M.C.N., Muotri, A.R., Griswold, M.D., et al. (2017) Intact piRNA pathway prevents L1 mobilization in male meiosis. Proc. Natl. Acad. Sci. U.S.A., 114.

7. Brouha, B., Schustak, J., Badge, R.M., Lutz-Prigge, S., Farley, A.H., Moran, J.V. and Kazazian, H.H. (2003) Hot L1s account for the bulk of retrotransposition in the human population. Proc. Natl. Acad. Sci. U.S.A., 100, 5280–5285.

8. Wildschutte, J.H., Williams, Z.H., Montesion, M., Subramanian, R.P., Kidd, J.M. and Coffin, J.M. (2016) Discovery of unfixed endogenous retrovirus insertions in diverse human populations. Proc Natl Acad Sci U S A, 113, E2326–2334.

9. Moran, J.V., DeBerardinis, R.J. and Kazazian, H.H. (1999) Exon shuffling by L1 retrotransposition. Science, 283, 1530–1534.

10. Xing, J., Wang, H., Belancio, V.P., Cordaux, R., Deininger, P.L. and Batzer, M.A. (2006) Emergence of primate genes by retrotransposon-mediated sequence transduction. Proc. Natl. Acad. Sci. U.S.A., 103, 17608–17613.

11. Belancio, V.P. (2006) LINE-1 RNA splicing and influences on mammalian gene expression. Nucleic Acids Research, 34, 1512–1521.

12. Raviram, R., Rocha, P.P., Luo, V.M., Swanzey, E., Miraldi, E.R., Chuong, E.B., Feschotte, C., Bonneau, R. and Skok, J.A. (2018) Analysis of 3D genomic interactions identifies candidate host genes that transposable elements potentially regulate. Genome Biol, 19, 216.

13. Zhang, Y., Li, T., Preissl, S., Amaral, M.L., Grinstein, J.D., Farah, E.N., Destici, E., Qiu, Y., Hu, R., Lee, A.Y., et al. (2019) Transcriptionally active HERV-H retrotransposons demarcate topologically associating domains in human pluripotent stem cells. Nat Genet, 51, 1380–1388.

14. Doronina, L., Reising, O., Clawson, H., Ray, D.A. and Schmitz, J. (2019) True Homoplasy of Retrotransposon Insertions in Primates. Syst Biol, 68, 482–493.

15. Sudmant, P.H., Rausch, T., Gardner, E.J., Handsaker, R.E., Abyzov, A., Huddleston, J., Zhang, Y., Ye, K., Jun, G., Hsi-Yang Fritz, M., et al. (2015) An integrated map of structural variation in 2,504 human genomes. Nature, 526, 75–81.

16. Dewannieux, M., Esnault, C. and Heidmann, T. (2003) LINE-mediated retrotransposition of marked Alu sequences. Nat Genet, 35, 41–48.

17. Payer, L.M. and Burns, K.H. (2019) Transposable elements in human genetic disease. Nat Rev Genet, 20, 760–772.

18. Kazazian, H.H., Wong, C., Youssoufian, H., Scott, A.F., Phillips, D.G. and Antonarakis, S.E. (1988) Haemophilia A resulting from de novo insertion of L1 sequences represents a novel mechanism for mutation in man. Nature, 332, 164–166.

19. Taşkesen, M., Collin, G.B., Evsikov, A.V., Güzel, A., Özgül, R.K., Marshall, J.D. and Naggert, J.K. (2012) Novel Alu retrotransposon insertion leading to Alström syndrome. Hum Genet, 131, 407–413.

20. Vogt, J., Bengesser, K., Claes, K.B.M., Wimmer, K., Mautner, V.-F., van Minkelen, R., Legius, E., Brems, H., Upadhyaya, M., Högel, J., et al. (2014) SVA retrotransposon insertion-associated deletion represents a novel mutational mechanism underlying large genomic copy number changes with non-recurrent breakpoints. Genome Biol, 15, R80.

21. Scott, E.C., Gardner, E.J., Masood, A., Chuang, N.T., Vertino, P.M. and Devine, S.E. (2016) A hot L1 retrotransposon evades somatic repression and initiates human colorectal cancer. Genome Res., 26, 745–755.

22. Upton, K.R., Gerhardt, D.J., Jesuadian, J.S., Richardson, S.R., Sánchez-Luque, F.J., Bodea, G.O., Ewing, A.D., Salvador-Palomeque, C., van der Knaap, M.S., Brennan, P.M., et al. (2015) Ubiquitous L1 mosaicism in hippocampal neurons. Cell, 161, 228–239.

23. Rodić, N., Steranka, J.P., Makohon-Moore, A., Moyer, A., Shen, P., Sharma, R., Kohutek, Z.A., Huang, C.R., Ahn, D., Mita, P., et al. (2015) Retrotransposon insertions in the clonal evolution of pancreatic ductal adenocarcinoma. Nat Med, 21, 1060–1064.

24. Hormozdiari, F., Hajirasouliha, I., Dao, P., Hach, F., Yorukoglu, D., Alkan, C., Eichler, E.E. and Sahinalp, S.C. (2010) Next-generation VariationHunter: combinatorial algorithms for transposon insertion discovery. Bioinformatics, 26, i350–357.

25. Quinlan, A.R., Clark, R.A., Sokolova, S., Leibowitz, M.L., Zhang, Y., Hurles, M.E., Mell, J.C. and Hall, I.M. (2010) Genome-wide mapping and assembly of structural variant breakpoints in the mouse genome. Genome Res, 20, 623–635.

26. Lee, E., Iskow, R., Yang, L., Gokcumen, O., Haseley, P., Luquette, L.J., Lohr, J.G., Harris, C.C., Ding, L., Wilson, R.K., et al. (2012) Landscape of somatic retrotransposition in human cancers. Science, 337, 967–971.

27. Keane, T.M., Wong, K. and Adams, D.J. (2013) RetroSeq: transposable element discovery from next-generation sequencing data. Bioinformatics, 29, 389–390.

28. Nakagome, M., Solovieva, E., Takahashi, A., Yasue, H., Hirochika, H. and Miyao, A. (2014) Transposon Insertion Finder (TIF): a novel program for detection of de novo transpositions of transposable elements. BMC Bioinformatics, 15, 71.

29. Thung, D.T., de Ligt, J., Vissers, L.E., Steehouwer, M., Kroon, M., de Vries, P., Slagboom, E.P., Ye, K., Veltman, J.A. and Hehir-Kwa, J.Y. (2014) Mobster: accurate detection of mobile element insertions in next generation sequencing data.

30. Wu, J., Lee, W.-P., Ward, A., Walker, J.A., Konkel, M.K., Batzer, M.A. and Marth, G.T. (2014) Tangram: a comprehensive toolbox for mobile element insertion detection. BMC Genomics, 15, 795.

31. Zhuang, J., Wang, J., Theurkauf, W. and Weng, Z. (2014) TEMP: a computational method for analyzing transposable element polymorphism in populations. Nucleic Acids Research, 42, 6826–6838.

32. Fiston-Lavier, A.-S., Barrón, M.G., Petrov, D.A. and González, J. (2015) T-lex2: genotyping, frequency estimation and re-annotation of transposable elements using single or pooled next-generation sequencing data. Nucleic Acids Res, 43, e22.

33. Jiang, C., Chen, C., Huang, Z., Liu, R. and Verdier, J. (2015) ITIS, a bioinformatics tool for accurate identification of transposon insertion sites using next-generation sequencing data. BMC Bioinformatics, 16, 72.

34. Chen, J., Wrightsman, T.R., Wessler, S.R. and Stajich, J.E. (2017) RelocaTE2: a high resolution transposable element insertion site mapping tool for population resequencing. PeerJ, 5, e2942.

35. Santander, C.G., Gambron, P., Marchi, E., Karamitros, T., Katzourakis, A. and Magiorkinis, G. (2017) STEAK: A specific tool for transposable elements and retrovirus detection in high-throughput sequencing data. Virus Evol, 3, vex023.

36. Gardner, E.J., Lam, V.K., Harris, D.N., Chuang, N.T., Scott, E.C., Pittard, W.S., Mills, R.E., The 1000 Genomes Project Consortium and Devine, S.E. (2017) The Mobile Element Locator Tool (MELT): population-scale mobile element discovery and biology. Genome Res., 27, 1916–1929.

37. Rajaby, R. and Sung, W.-K. (2018) TranSurVeyor: an improved database-free algorithm for finding non-reference transpositions in high-throughput sequencing data. Nucleic Acids Res, 46, e122.

38. Chen, X. and Li, D. (2019) ERVcaller: identifying polymorphic endogenous retrovirus and other transposable element insertions using whole-genome sequencing data. Bioinformatics, 35, 3913–3922.

39. Sanchez-Luque, F.J., Kempen, M.-J.H.C., Gerdes, P., Vargas-Landin, D.B., Richardson, S.R., Troskie, R.-L., Jesuadian, J.S., Cheetham, S.W., Carreira, P.E., Salvador-Palomeque, C., et al. (2019) LINE-1 Evasion of Epigenetic Repression in Humans. Mol Cell, 75, 590-604.e12.

40. Puurand, T., Kukuškina, V., Pajuste, F.-D. and Remm, M. (2019) AluMine: alignment-free method for the discovery of polymorphic Alu element insertions. Mob DNA, 10, 31.

41. Goubert, C., Thomas, J., Payer, L.M., Kidd, J.M., Feusier, J., Watkins, W.S., Burns, K.H., Jorde, L.B. and Feschotte, C. (2020) TypeTE: a tool to genotype mobile element insertions from whole genome resequencing data. Nucleic Acids Research, 48, e36–e36.

42. Chu, C., Borges-Monroy, R., Viswanadham, V.V., Lee, S., Li, H., Lee, E.A. and Park, P.J. (2021) Comprehensive identification of transposable element insertions using multiple sequencing technologies. Nat Commun, 12, 3836.

43. Eraslan, G., Avsec, Ž., Gagneur, J. and Theis, F.J. (2019) Deep learning: new computational modelling techniques for genomics. Nat Rev Genet, 20, 389–403.

44. Jaganathan, K., Kyriazopoulou Panagiotopoulou, S., McRae, J.F., Darbandi, S.F., Knowles, D., Li, Y.I., Kosmicki, J.A., Arbelaez, J., Cui, W., Schwartz, G.B., et al. (2019) Predicting Splicing from Primary Sequence with Deep Learning. Cell, 176, 535-548.e24.

45. Zhang, Y., An, L., Xu, J., Zhang, B., Zheng, W.J., Hu, M., Tang, J. and Yue, F. (2018) Enhancing Hi-C data resolution with deep convolutional neural network HiCPlus. Nat Commun, 9, 750.

46. Wang, J., Lisanza, S., Juergens, D., Tischer, D., Watson, J.L., Castro, K.M., Ragotte, R., Saragovi, A., Milles, L.F., Baek, M., et al. (2022) Scaffolding protein functional sites using deep learning. Science, 377, 387–394.

47. Poplin, R., Chang, P.-C., Alexander, D., Schwartz, S., Colthurst, T., Ku, A., Newburger, D., Dijamco, J., Nguyen, N., Afshar, P.T., et al. (2018) A universal SNP and small-indel variant caller using deep neural networks. nature biotechnology, 36, 9.

48. Bao, W., Kojima, K.K. and Kohany, O. (2015) Repbase Update, a database of repetitive elements in eukaryotic genomes. Mob DNA, 6, 11.

49. Byrska-Bishop, M., Evani, U.S., Zhao, X., Basile, A.O., Abel, H.J., Regier, A.A., Corvelo, A., Clarke, W.E., Musunuri, R., Nagulapalli, K., et al. (2022) High-coverage whole-genome sequencing of the expanded 1000 Genomes Project cohort including 602 trios. Cell, 185, 3426-3440.e19.

50. Robinson, J.T., Thorvaldsdóttir, H., Winckler, W., Guttman, M., Lander, E.S., Getz, G. and Mesirov, J.P. (2011) Integrative genomics viewer. Nat Biotechnol, 29, 24–26.

51. Szegedy, C., Vanhoucke, V., Ioffe, S., Shlens, J. and Wojna, Z. (2016) Rethinking the Inception Architecture for Computer Vision. In 2016 IEEE Conference on Computer Vision and Pattern Recognition (CVPR). IEEE, Las Vegas, NV, USA, pp. 2818–2826.

52. Martín Abadi, Ashish Agarwal, Paul Barham, Eugene Brevdo, Zhifeng Chen, Craig Citro, Greg S. Corrado, Andy Davis, Jeffrey Dean, Matthieu Devin, et al. (2015) TensorFlow: Large-Scale Machine Learning on Heterogeneous Systems.

53. Li, H. and Durbin, R. (2009) Fast and accurate short read alignment with Burrows-Wheeler transform. Bioinformatics, 25, 1754–1760.

54. Payer, L.M., Steranka, J.P., Yang, W.R., Kryatova, M., Medabalimi, S., Ardeljan, D., Liu, C., Boeke, J.D., Avramopoulos, D. and Burns, K.H. (2017) Structural variants caused by Alu insertions are associated with risks for many human diseases. Proc. Natl. Acad. Sci. U.S.A., 114.

55. Hinrichs, A.S., Karolchik, D., Baertsch, R., Barber, G.P., Bejerano, G., Clawson, H., Diekhans, M., Furey, T.S., Harte, R.A., Hsu, F., et al. (2006) The UCSC Genome Browser Database: update 2006. Nucleic Acids Res, 34, D590–598.

56. McLaren, W., Gil, L., Hunt, S.E., Riat, H.S., Ritchie, G.R.S., Thormann, A., Flicek, P. and Cunningham, F. (2016) The Ensembl Variant Effect Predictor. Genome Biol, 17, 122.

57. Bennett, E.A., Keller, H., Mills, R.E., Schmidt, S., Moran, J.V., Weichenrieder, O. and Devine, S.E. (2008) Active Alu retrotransposons in the human genome. Genome Res, 18, 1875–1883.

58. Derrien, T., Estellé, J., Marco Sola, S., Knowles, D.G., Raineri, E., Guigó, R. and Ribeca, P. (2012) Fast computation and applications of genome mappability. PLoS One, 7, e30377.

59. Larson, P.A., Moldovan, J.B., Jasti, N., Kidd, J.M., Beck, C.R. and Moran, J.V. (2018) Spliced integrated retrotransposed element (SpIRE) formation in the human genome. PLoS Biol, 16, e2003067.

60. Watkins, W.S., Feusier, J.E., Thomas, J., Goubert, C., Mallick, S. and Jorde, L.B. (2020) The Simons Genome Diversity Project: A Global Analysis of Mobile Element Diversity. Genome Biology and Evolution, 12, 779–794.

61. Collins, R.L., Brand, H., Karczewski, K.J., Zhao, X., Alföldi, J., Francioli, L.C., Khera, A.V., Lowther, C., Gauthier, L.D., Wang, H., et al. (2020) A structural variation reference for medical and population genetics. Nature, 581, 444–451.

