## Supplemental Figure S1-9 for "Identification of mobile element insertion from whole genome sequencing data using deep neural network model"

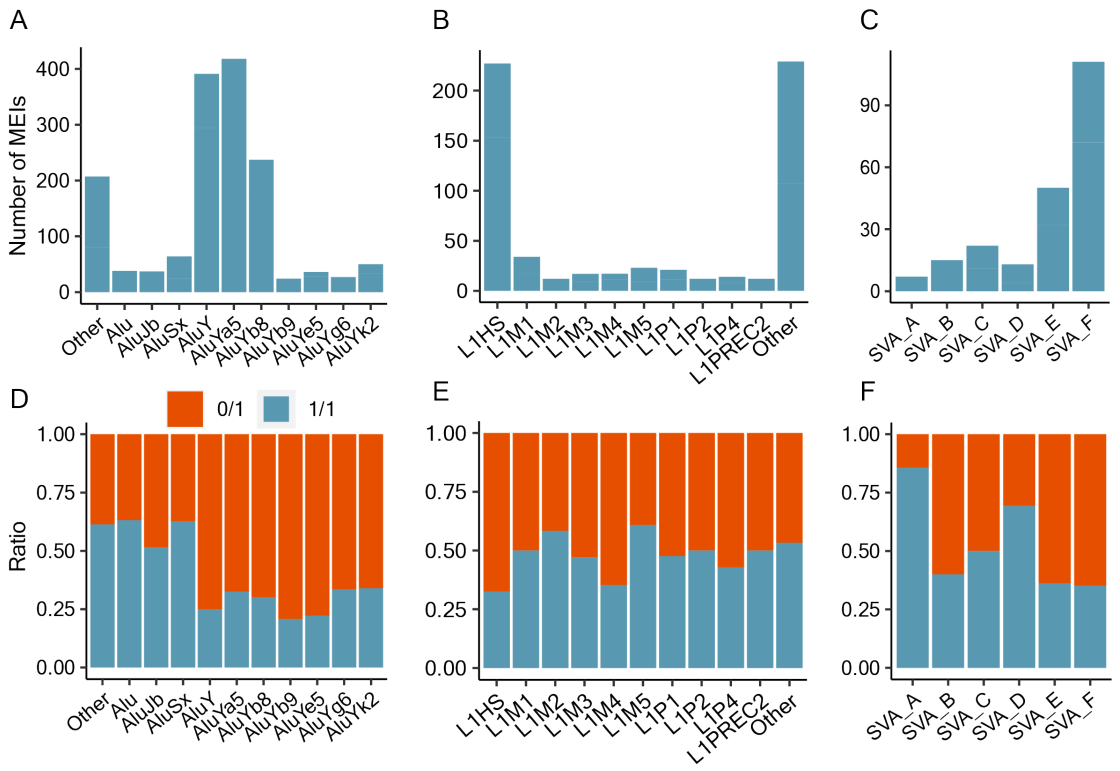


**Figure S1.** Count of non-reference ME types in the GIAB HG002 benchmark dataset. Number of **A)** Alu, **B)** L1, and **C)** SVA insertions by subtype. Homozygous and heterozygous ratio of **D)** Alu, **E)** L1, and **F)** SVA insertions by subtype.

**Figure S2**

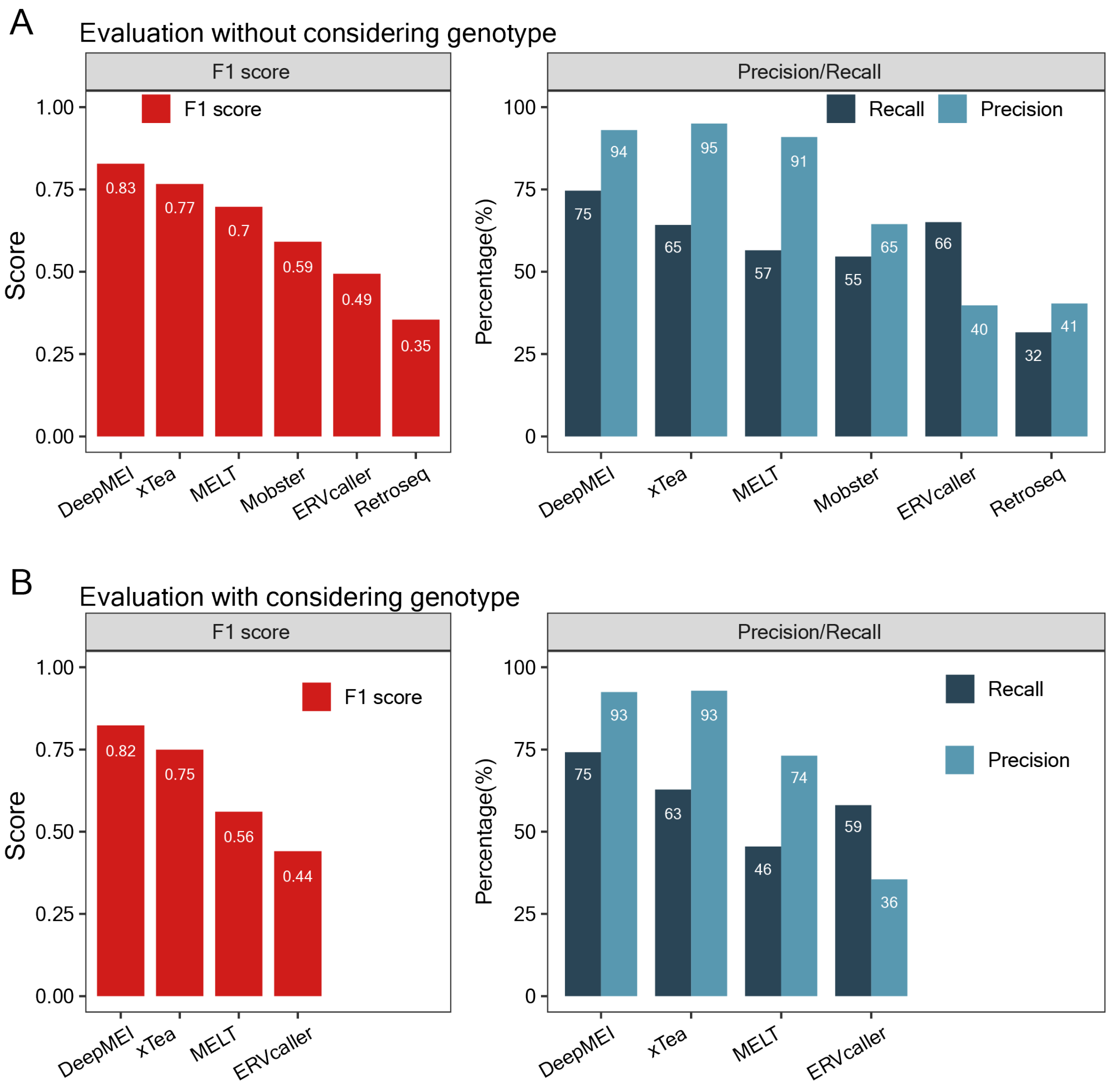


**Figure S2.** Performance comparison of DeepMEI with five existing tools for analyzing Alu insertions in the GIAB HG002 benchmark dataset. **A)** The first evaluation was based on the called Alu sites without considering genotype information. **B)** The second evaluation was based on the called Alu sites considering correct genotype information. Genotype output was unavailable in Mobster and Retroseq, and the two tools were not included in the genotype-based evaluation.

**Figure S3**

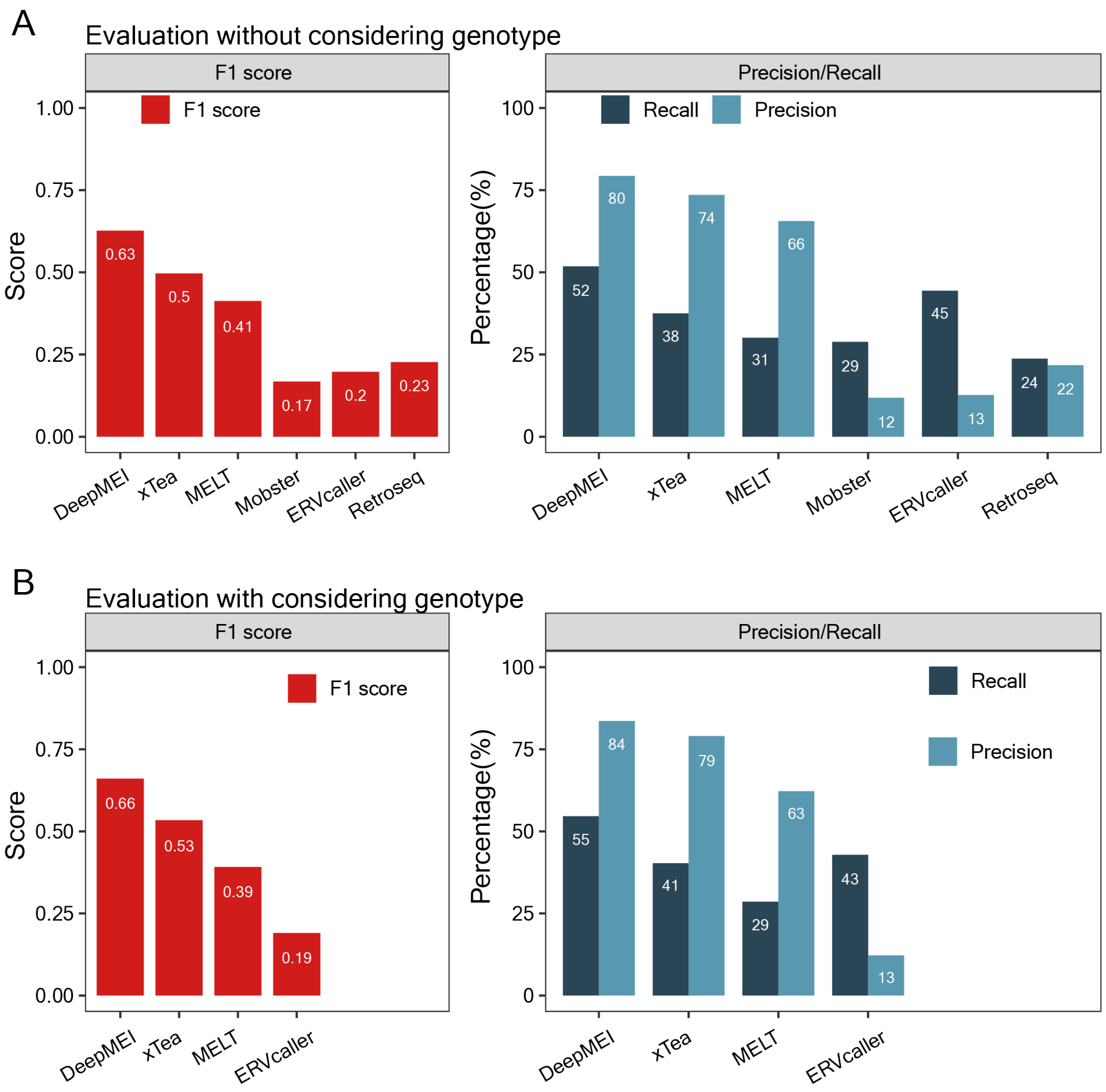


**Figure S3.** Performance comparison of DeepMEI with five existing tools for analyzing L1 insertions in the GIAB HG002 benchmark dataset. **A)** The first evaluation was based on the called L1 sites without considering genotype information. **B)** The second evaluation was based on the called L1 sites considering correct genotype information. Genotype output was unavailable in Mobster and Retroseq, and the two tools were not included in the genotype-based evaluation.

**Figure S4**

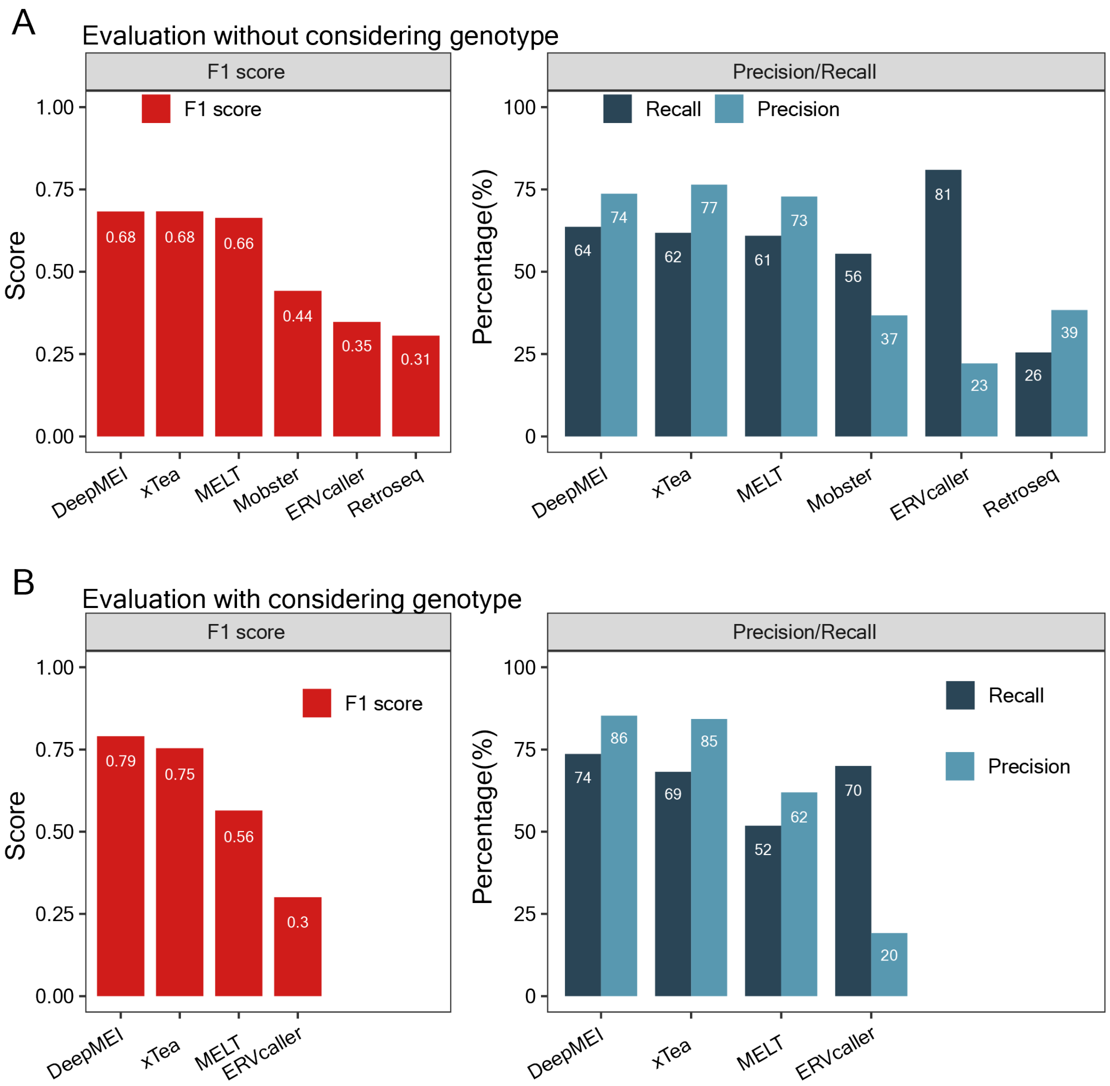


**Figure S4.** Performance comparison of DeepMEI with five existing tools for analyzing SVA insertions in the GIAB HG002 benchmark dataset. **A)** The first evaluation was based on the called SVA sites without considering genotype information. **B)** The second evaluation was based on the called SVA sites considering correct genotype information. Genotype output was unavailable in Mobster and Retroseq, and the two tools were not included in the genotype-based evaluation.

**Figure S5**

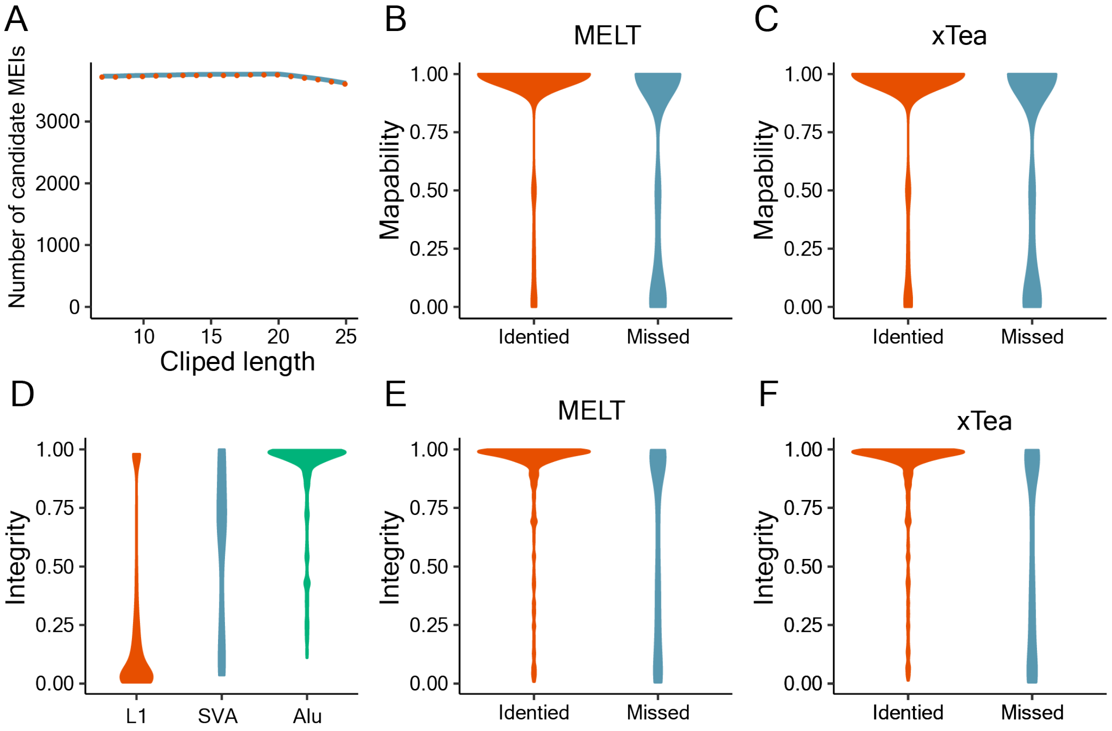


**Figure S5.** Evaluation of the factors affecting MEI identification. **A)** correlation between the number of candidate MEI sites and clipped length of reads. **B)** Low genome mappability reduced the performance of MELT. **C)** Low genome mappability reduced the performance of xTea. **D)** Integrity distribution of Alu, L1, and SVA in the GIAB HG002 benchmark dataset. Most L1 insertions were truncated. **E)** Low ME integrity reduced the performance of MELT. **F)** Low ME integrity reduced the performance of xTea.

**Figure S6**

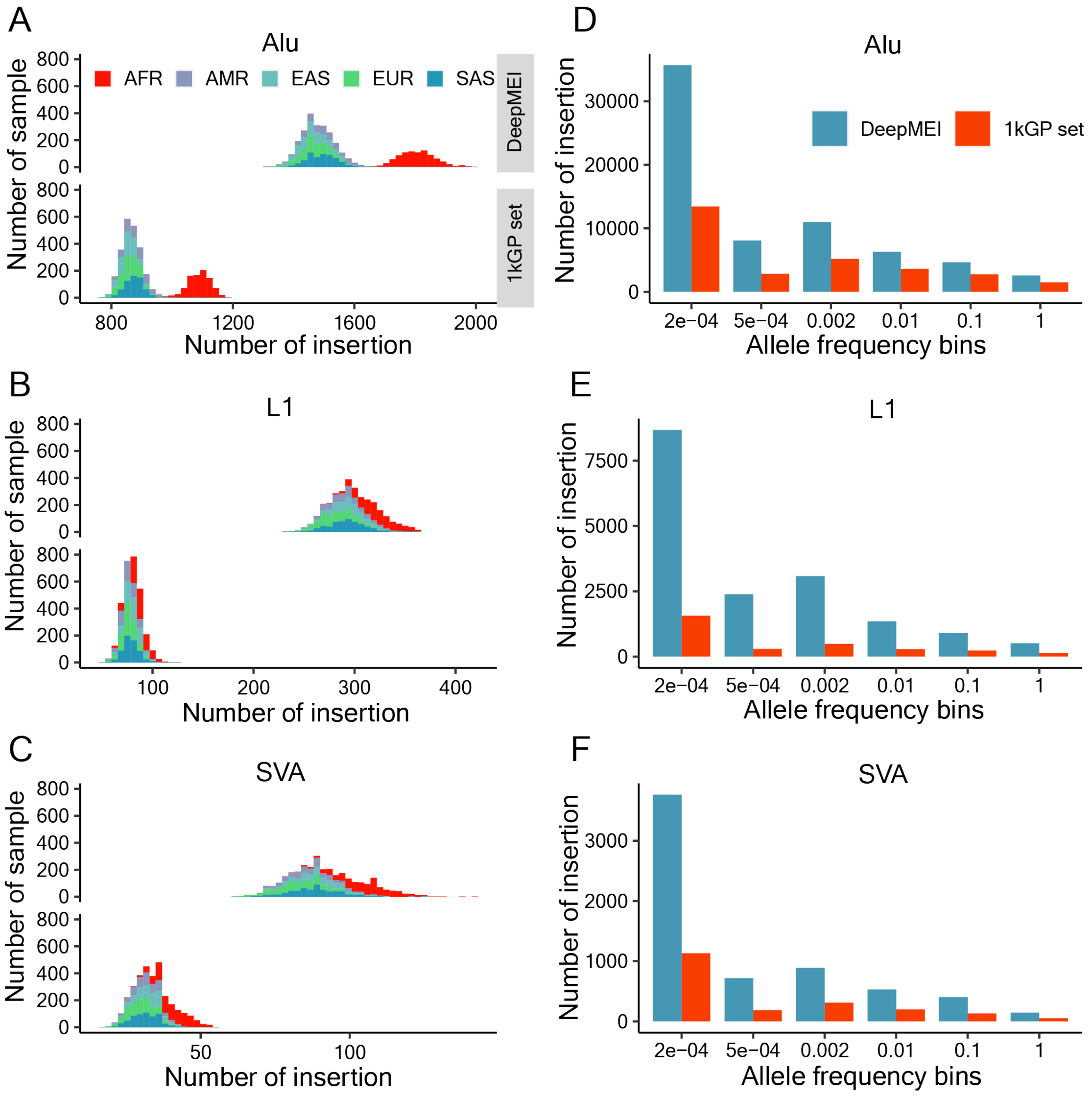


**Figure S6.** Distribution of MEI number per sample and allele frequency. Population abbreviations are AFR: Africa, AMR: America, EAS: East Asia, EUR: Europe, SAS: South Asia.

**Figure S7**

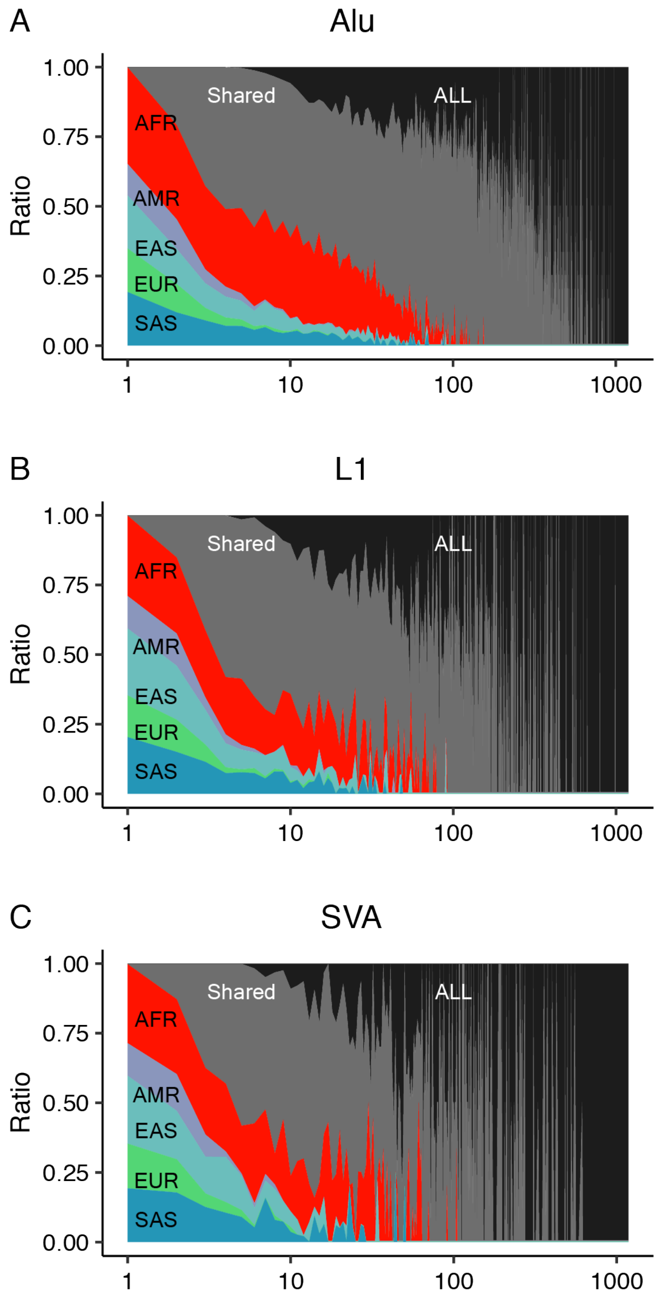


**Figure S7.** Unique and shared non-reference Alu, L1, and SVA insertions identified in data from different populations from 1kGP.

**Figure S8**

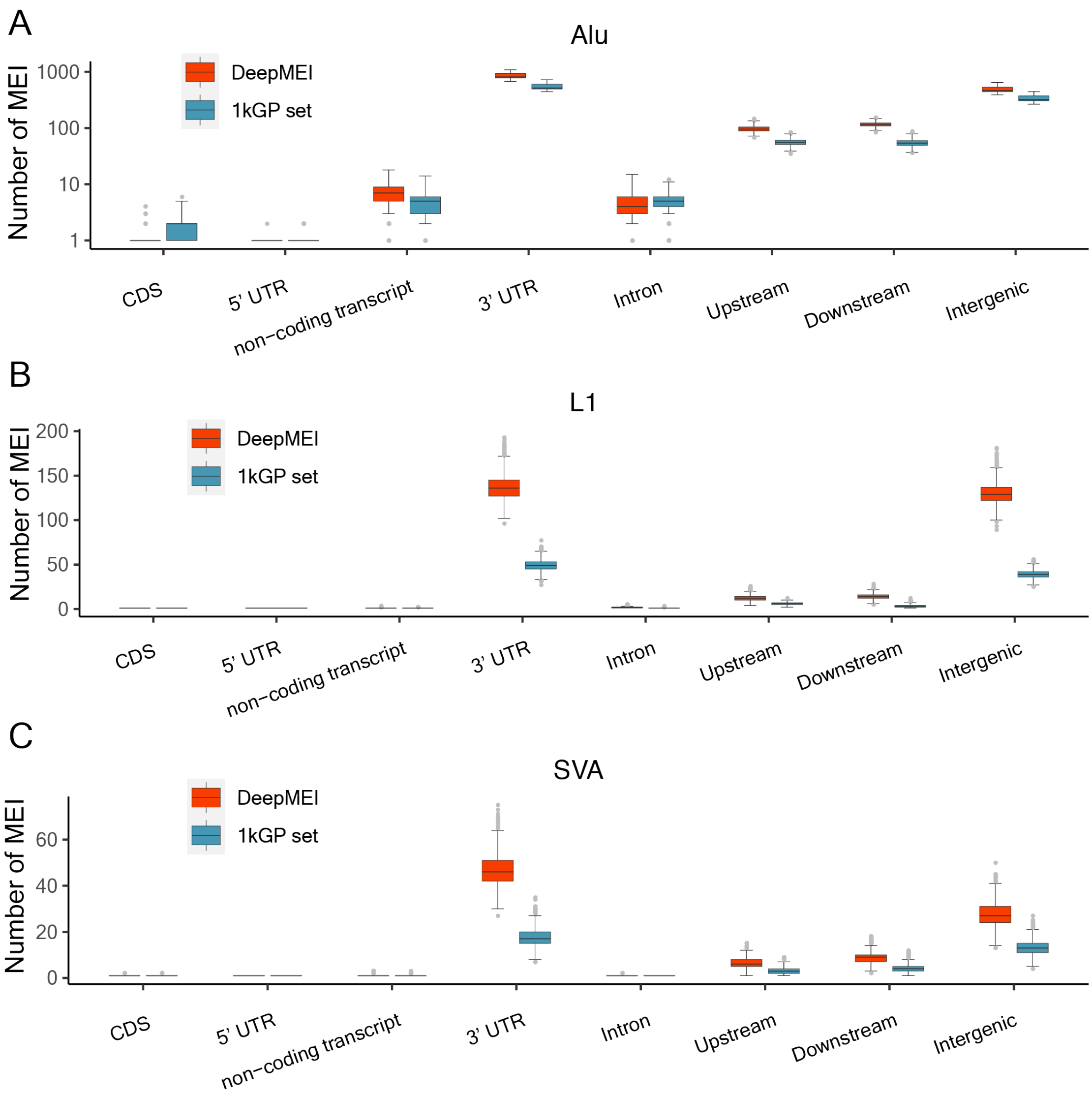


**Figure S8.** Distributions of non-reference ME count and enrichment analysis by genome region

**Figure S9**

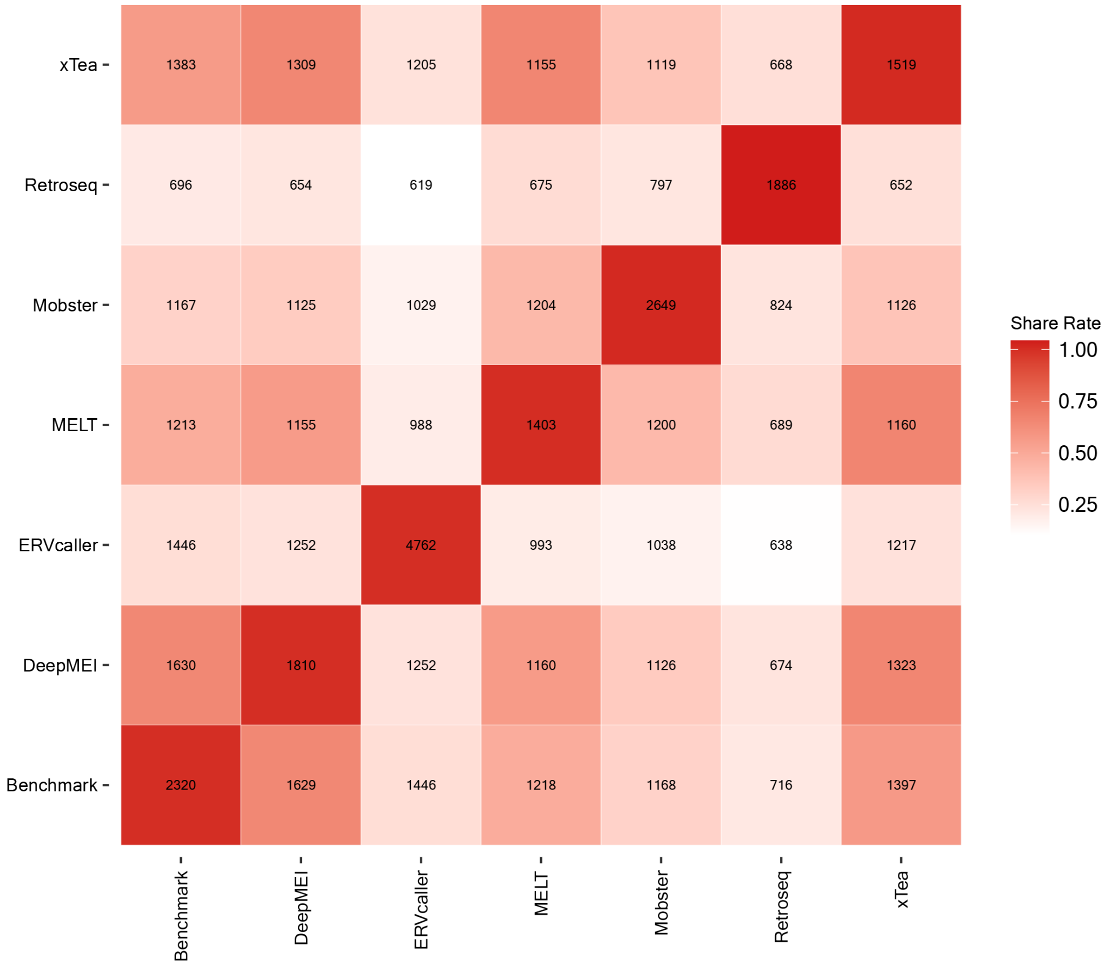


**Figure S9.** Overlapping MEIs between pairs of tools
